# Noninvasive ultrasound targeted modulation of calcium influx in splenic immunocytes potentiates antineoplastic immunity attenuating hepatocellular carcinoma proliferation

**DOI:** 10.1101/2025.03.31.646454

**Authors:** Wei Dong, Guihu Wang, Senyang Li, Yichao Chai, Qian Wang, Yucheng Li, Qiaoman Fei, Yujin Zong, Jing Geng, Pengfei Liu, Zongfang Li

**Author notes:** **Correspondence authors:** Zongfang Li,; Pengfei Liu. First author: Wei Dong,.

## Abstract

The spleen, as the largest immune organ, plays a pivotal role in modulating immune responses, particularly in the context of carcinogenesis and tumor progression. Non-pharmacological manipulation, particularly splenic ultrasound stimulation (SUS), has demonstrated significant immunomodulatory efficacy in alleviating chronic inflammatory diseases, suggesting its potential to revitalize splenic immunocompetence suppressing tumor proliferation, yet remains underexplored. This study applied low-frequency pulsed focused ultrasound (FUS) noninvasively stimulating the spleen (FUS sti. spleen) to investigate the efficacy in enhancing antitumor immunity and suppressing hepatocellular carcinoma (HCC). The results showed that FUS sti. spleen significantly suppressed tumor proliferation, achieving a suppression rate of >70% for H22-HCC and >83% for Hepa1-6-HCC, along with significantly prolonged survival. Comprehensive flow cytometry, single-cell RNA sequencing (scRNA-seq) and cytokine analyses demonstrated that SUS profoundly reshaped the splenic and intratumoral immune landscape, specifically activating cytotoxic CD8^+^ T cells and NK cells while suppressing immunosuppressive cell populations. Mechanistically, FUS facilitated calcium influx in splenic immunocytes, activating multiple signaling pathways, such as TNF, NFκB, MAPK, HIF-1, and ErbB, thereby counteracting tumor-driven immunosuppressive polarization while potentiating robust immune activation that impedes malignant progression and neoplastic proliferation. Leveraging above insights, we developed spleen-targeted nanodroplets encapsulating bioavailable calcium ions (STNDs@Ca²⁺), which, upon FUS stimulation, undergo cavitation-mediated controlled release of Ca²⁺, further amplifying immunocyte activation and tumor suppression, achieving a remarkable H22-HCC suppression rate of over 90%. This study highlights the therapeutic potential of ultrasound-mediated splenic immunomodulation, both as a standalone intervention and in synergy with STNDs@Ca²⁺, as a novel and noninvasive strategy for cancer immunotherapy.

## 1 Introduction

Spleen is the largest immune organ correlated to the occurrence and development of many diseases that attracted great attention and in-depth study by researchers and clinicians since the report of a huge post-splenectomy infection after splenectomy in the 1930s [1–3]. In particular, the splenic immunity was closely related to carcinogenesis and tumor progression, in which the spleen exerted anti-tumor effects in the early stage [4], but induced an “inhibitory” immune property that promoted tumor growth, invasion, and metastasis in the middle-late stage [5]. Accordingly, the above experimental results were replicated using an *in situ* xenograft hepatoma-22 hepatocellular carcinoma (H22-HCC) model, in which the proportion of CD3^+^/CD4^+^ T cells significantly increased during the first week, but synchronized with a significant decrease of CD8 T cells and NK cells at the second week. This was followed with the proportion of myeloid-derived suppressor cells (MDSCs) and Treg cells showing an evident increase after two weeks [6]. Thus, it was therefore investigated whether non-pharmacological manipulation of splenic immunity could be used to treat malignancies.

Despite the lack of relevant literature on the treatment of tumors, in the last decade, numerous studies have been performed on the therapy of inflammatory diseases by directly or indirectly modulating the immune property of the spleen through physical means such as electrodes and ultrasound. The implanted or transcutaneous electrodes stimulating spleen have been used to improve inflammatory disease (i.e., rheumatoid arthritis [7], hypertension [8], sepsis [9], kidney ischemia-reperfusion injury [10], inflammatory bowel disease [11], and systemic lupus erythematosus (clinical trial: NCT02822989)) mainly by modulating the cholinergic anti-inflammatory pathway represented by the vagus nerve and the spleen circuit [12]. However, the transcutaneous electrodes stimulate the superficial organs and tissues with non-targeted performance, and even the invasively implanted electrodes can only stimulate a single main nerve in a limited space, thus making difficult the accurate intervention on the spleen, causing certain side effects, including the reduction of heart rate [13, 14].

Interestingly, focused ultrasound (FUS), as another noninvasive therapeutic option, highlighted the potential feasibility to alleviate chronic inflammatory diseases with equivalent efficacy to electrodes-based vagus nerve stimulation, but possessing more significant advantages such as deep tissue penetration, precise targeting, and manipulability [15–18]. Accordingly, Okusa et al. [19], Cotero et al. [20], Zachs et al. [14], and others [21] have empirically validated the therapeutic efficacy of FUS modulating splenic immunologic function in the management of diverse pathological conditions, encompassing rheumatoid arthritis, colitis, pneumonia, hyperglycemia, and other inflammatory disorders. Their seminal investigations have further elucidated that the underlying mechanism of splenic ultrasound stimulation (SUS) predominantly involves the activation of the cholinergic anti-inflammatory pathway (CAP), providing crucial insights into the splenic neuroimmunological basis [20, 22, 23]. Irrationally, the majority of research has focused on the SUS modulation of inflammatory mediators, such as TNF-α, IL-6, and IL-10, as well as on CAP validation, but exists a conspicuous paucity of studies elucidating the dynamic phenotypic and functional alterations in diverse splenocyte populations during SUS. Concurrently, emerging evidence has substantiated that ultrasound irradiation exerts direct modulatory effects on immunocyte immunocompetence through multifaceted biophysical mechanisms, including the transcriptional regulation of immunologically relevant genes and signaling cascades, perturbation of cytokine networks, and facilitation of transmembrane ion flux [14, 24–26]. Besides, they highlighted that ultrasound potentiated cellular proliferation, activation, migration, infiltration, and phagocytic capacity, mainly attributed to ultrasonic mechanical forces, microjets, vaporization, and cavitation effects within the splenic parenchyma. Additionally, several studies have reported significant alterations in the proportions of T cells and B cells, but not Mφs (the principal executor of CAP), in the spleen following SUS, highlighting the importance to study the direct modulation of immune cells by ultrasound [24, 27]. Therefore, the dependence, specificity, and mechanism of spleen responding to ultrasonic stimulation need to be scientifically and clinically verified in a larger cohort. Of paramount importance is the comprehensive profiling of the heterogeneous immunophenotypic and functional responses across distinct splenic lymphocyte lineages and subsets to ultrasonic modulation, with special emphasis on elucidating the dynamic transition of specialized immunocyte populations that exhibit unique mechanosensitivity to acoustic stimuli. Collectively, these findings herald the advent of ultrasound-mediated splenic immunomodulation as a transformative paradigm in cancer immunotherapy, necessitating immediate and systematic investigation to elucidate its therapeutic potential and mechanistic underpinnings.

This study applied low-frequency pulsed FUS noninvasively stimulating the spleen (FUS sti. spleen) to investigate the curative effect of HCC suppression (such as orthotopic H22-HCC and Hepa1-6-HCC), and performed flow cytometry (FCM) to detect immunocyte heterogeneity in the spleen, peripheral blood, tumor, and para-carcinoma tissues. Additionally, the single-cell RNA sequencing (scRNA-seq) was applied to delineate the splenic and intratumoral multifaceted immune landscape, dynamic transition, and interactome underlying the therapeutic FUS sti. spleen. Of particular significance, CD8 T lymphocytes and NK cells emerged as the predominant cellular subsets demonstrating remarkable responsiveness to FUS intervention. Mechanistic investigations further demonstrated that ultrasound-facilitated Ca^2+^ influx orchestrates a cascade of intracellular signaling pathways, such as TNF, NFκB, MAPK, HIF-1, calcium and ErbB, culminating in the activation of splenic antitumor immunity and subsequent suppression of HCC proliferation. Capitalizing on these findings, we engineered spleen-targeted nanodroplets encapsulating bioavailable calcium ions (STNDs@Ca^2+^), which, upon ultrasound stimulation, undergo cavitation-mediated controlled release of exogenous Ca^2+^. Simultaneously inducing sonoporation-mediated enhancement of membrane permeability facilitates cellular Ca^2+^ uptake, thereby potentiating the tumoricidal capacity of splenic immunocytes at elevated calcium concentrations to activate calcium-related signaling pathways, ultimately achieving superior tumor suppression.

## 2 Materials and methods

### 2.1 Cell line and animal model

H22 and Hepa1-6 cell lines were purchased from the Cell Bank of the Chinese Academy of Sciences (Shanghai, China). H22 cells were multiplied in ascites after intraperitoneal injection in mice (approximately 1.0*10^7 cells/mL; 200 µL per mouse). Hepa1-6 cells cells were cultured in Dulbecco’s modified Eagle’s medium (Cat. No. 11320033, Gibco, Invitrogen, Carlsbag, USA) supplemented with 10% fetal bovine serum (FBS; Cat. No. A5669701, Gibco, Invitrogen, Carlsbag, USA) and 1% penicillin-streptomycin solution (Cat. No. 15140122, Gibco, Invitrogen, Carlsbag, USA), and incubated at 37 °C under 5% CO2 and 100% humidity. The *in situ* xenograft HCC model was created by an incision of approximately 0.8 cm in length on the upper end of the midabdominal line after the mouse was anesthetized with 1.5% isohalothane and fixed on the anatomic stage. A total of 25 µL H22/Hepa1-6 cells (approximately 1.0*10^7 cells/mL) were injected into the left hepatic lobe using an insulin syringe, and the incision was closed after applying pressure on the pinhole with a medical cotton swab for 3-4 minutes. The mice used in this experiment were 7-week male C57black/6 weighting 16.5-17.5 g and obtained from the Beijing Animal Experiment Center (Chinese Academy of Sciences, Beijing, China), and were acclimatized for at least 1 week before the experiments.

The tumor volume was calculated as follows: volume = 0.5*L*W^2, where L and W were the length and width of the tumor, respectively, measured by a caliper. The weight of the spleen, liver, tumor, and mice was assessed by electronic scales, and the amount of ascites was quantified using a syringe.

### 2.2 Ultrasonic platform applied for FUS sti. spleen

The diagram of ultrasonic platform in Figure 1 A shows a function generator (Cat. No. 33120A, Agilent, Santa Clara, USA) that produced a pulsed sinusoidal waveform triggering the power amplifier (Cat. No. AG1006, California, USA) to drive a 1.04 MHz FUS transducer (Chongqing Ronghai Engineering Research Center of Ultrasound Medicine Co., Ltd, Chongqing, China) with a 100 mm aperture and 65 mm focus, whose focal region was approximately 1.4*1.4*8.6 mm^3^. The FUS transducer was mounted on a XYZ motorized positioning stage to control SUS periods, and the position of the FUS focal region relative to spleen was adjusted under a B-mode imaging guidance with a 3.2 MHz phased array embedded into the center opening of the FUS transducer. The mice were anesthetized with 1.5% inhaled isoflurane and placed on a manual translation stage equipped with a heating pad. Then, a centrifuged coupling gel was immediately applied to the shaved skin, and the manual translation stage was adjusted to allow the tight contact of the spleen region with the bottom of the water tank under the B-mode imaging guidance. The FUS was moved across the whole spleen through the XYZ programmable logical controller, which was performed every two days for 25 minutes (Figure 1 B). The acoustic pressure and spatial beam profile of the FUS transducer (parameters set at 2.4 MPa peak negative pressure and 10% duty cycle) were calibrated by a hydrophone (Cat. No. HNR-1000, ONDA, Videlles, France) before performing SUS (Figure 1 C).

**Figure 1.**
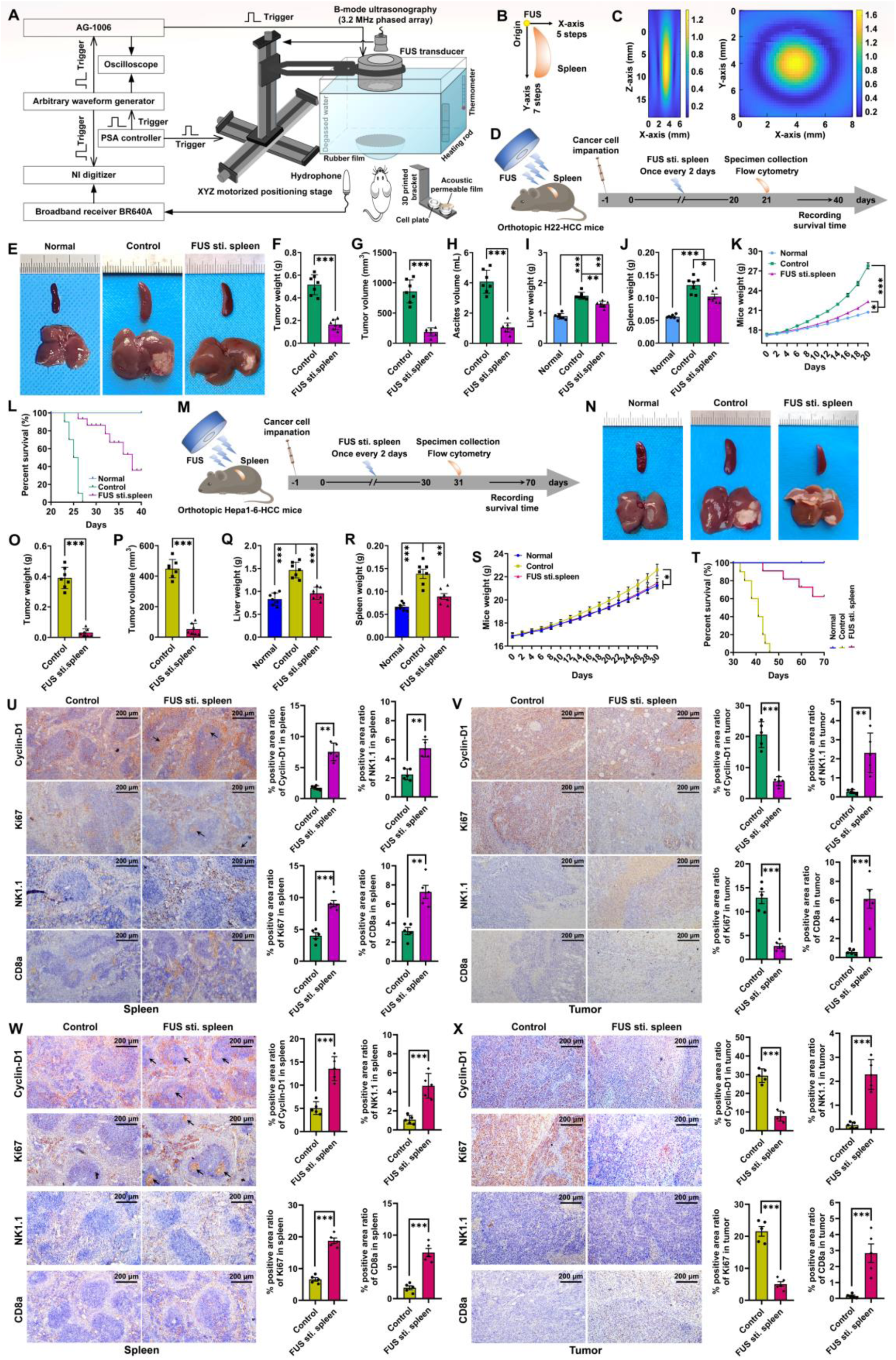
FUS sti. spleen suppressed orthotopic H22/Hepa1-6-HCC proliferation. A, the diagram of ultrasonic platform applied for FUS sti. spleen. B, the scheme of whole spleen irradiated by FUS transducer carried by the XYZ motorized positioning stage. C, ultrasound focal field scanning after calibration by a hydrophone. D, experimental protocol for the use of FUS sti. spleen after H22 cancer cell implantation into the liver. E, images of spleen and liver bearing H22-HCC tumor. F-G, statistics of H22-HCC tumor weight and volume, respectively. H, H22 malignant ascites volume. I-K, liver weight (total weight of liver parenchyma and H22-HCC tumor), spleen weight, and mice weight, respectively. L, Kaplan-Meier survival curves of G1-G3 groups. M, schematic diagram of the experimental protocol for FUS sti. spleen following orthotopic implantation of Hepa1-6 cancer cells in the liver. N, representative images showing the spleen and liver with Hepa1-6-HCC tumor. O-P, statistics of Hepa1-6-HCC tumor weight and volume, respectively. Q-S, evaluation of total liver weight (including both parenchyma and Hepa1-6-HCC tumor), spleen weight, and body weight, respectively. T, Kaplan-Meier survival curves of T1-T3 groups. U-V, immunohistochemical staining of cyclin-D1, Ki67, NK1.1, and CD8a positive cells in the spleen and tumor from G2-G3 groups, and the corresponding quantification of positive area ratio, respectively. W-X, immunohistochemical staining and corresponding quantitative analysis of cyclin-D1, Ki67, NK1.1, and CD8a expression in both splenic and tumor tissues from T2-T3 groups. G1 and T1 group with normal mice classified as negative control (named normal), G2 group with orthotopic H22-HCC mice and T2 group with orthotopic Hepa1-6-HCC mice used as the positive control (named control), and G3 group with orthotopic H22-HCC mice and T3 group with orthotopic Hepa1-6-HCC mice respectively subjected to FUS stimulation (named FUS sti. spleen).

### 2.3 Flow cytometry

The cell suspensions from the spleen, peripheral blood, tumor, and para-carcinoma tissue were filtrated through a 70-μm nylon cell strainer, washed with sterilized PBS, and incubated with the antibodies listed in Supplementary Table 1. The scheme of antibody labeling for each type of immune cell is shown in Supplementary Table 2. The immune cells were counted by FCM using CytoFLEX LX (Beckman Coulter Life Sciences, USA), and analyzed using the FlowJo software (FlowJo 10, LLC, Ashland, OR).

To count Ca^2+^ positive cells, splenic cell suspensions were incubated with Fluo-4 AM dye (1 mL dye/10^6^ cells; Cat. No. S1061M, Beyotime, Shanghai, China) for 30 min at 37 ℃ according to the manufacturer’s instructions. The fluorescent cells were counted by FCM using the CytoFLEX LX (Ex/Em = 490/525 nm).

### 2.4 Histological and immunohistochemical staining

The tissues were cut into 5 μm-thick sections prepared by a routine protocol, followed by incubation with primary antibodies against cyclin-D1 (Cat. No. ab273610, Abcam, Cambridge, UK), Ki67 (Cat. No. RM9106S1, Thermo Fisher Scientific, MA, USA), NK1.1 (Cat. No. 108759, BioLegend, California, USA), and CD8a (Cat. No. ab4055, Abcam, Cambridge, UK) for immunohistochemical staining to verify immune cell alteration due to SUS. To detect calcium deposits (mineralization), the tissue sections were stained using the VON KOSSA Calcium Staining Kit (Cat. No. JM1519, HPBIO, Shanghai, China), where mass deposits appeared black and dispersed deposits appeared gray. Additionally, to assess tissue damage, the histologic sections were stained with hematoxylin-eosin (HE) and TdT-dependent dUTP-biotin nick end labeling (TUNEL) staining. Images were captured in at least 5-10 different regions randomly selected in each section using a fluorescence microscopy (observer3, Carl Zeiss, Jena, Germany), and the positive area ratio was quantified using the Image Pro Plus 6.0 software (Media Cybernetics, CA, United States).

### 2.5 scRNA-seq, scTCR-seq and data processing

The splenic and intratumoral CD45^+^ cells were isolated via the ultra-high speed FCM sorting system (MoFlo Astrios EQ, Beckman Coulter, lnc., USA). Particularly, the sorted intratumoral CD45^+^ cells need to be mixed with the original tumor cell suspension at a ratio of 3 : 1. Single-cell libraries were constructed using the 10× Genomics Chromium platform, followed by sequencing on the Illumina Novaseq 6000. Data processing involved alignment to the mouse reference genome (mm10) using Cell Ranger software (version 7.0.1), with quality control, normalization, and batch effect correction performed using Seurat [28] and batchelor packages [29]. Pseudotime trajectory analysis was conducted with Monocle2 to elucidate dynamic immune cell transitions [30], while Cell Chat R package (version 1.1.3) was employed to map intercellular communication networks. Copy number variation (CNV) analysis using the inferCNV package identified malignant cells [31]. TCR sequencing data were analyzed with the Python toolkit scirpy (version 0.11.1), focusing on clonotype identification and clonality assessment. The comprehensive details are provided in Supplementary File 1.

### 2.6 bRNA-seq and GSEA analysis

The splenic CD8 T cells (CD45^+^ CD3^+^ CD8a^+^) and NK cells (CD45^+^ CD3^-^ NK1.1^+^) were sorted from the FUS sti. spleen group and the control group, respectively, using the Beckman Kurt MoFlo Astrios ultra high-speed FCM sorting system. The total RNA was extracted using TRIZOL (Cat. No. 15596026CN, Thermo Fisher Scientific, MA, USA), reverse transcribed to cDNA using the SuperScript IV CellsDirect cDNA Synthesis Kit (Cat. No. 11750150, Thermo Fisher Scientific, MA, USA), and subjected to bulk RNA sequencing (bRNA-seq) by the OE Biotech Co., Ltd. (Shanghai, China). Gene set enrichment analysis (GSEA) was performed using the clusterProfiler package for all genes in the samples of the FUS sti. spleen group and the control group in TCGA to explore the difference in function and associated pathways between both groups.

### 2.7 qRT-PCR

qRT-PCR was performed using the SuperScript™ III Platinum™ One-Step qRT-PCR Kit (Cat. No. 11732088, Thermo Fisher Scientific, MA, USA) and the 7500 Real-Time PCR System (Applied Biosystems, Foster City, CA, USA). All primer sequences are listed in Supplementary Table 3.

### 2.8 Calcium-related *in vitro* experiments

The CD8 T cells (CD45^+^ CD3^+^ CD8a^+^) and NK cells (CD45^+^ CD3^-^ NK1.1^+^) were sorted and subjected to FUS stimulation *in vitro* with or without high calcium concentration (300 nM). Cancer cell lysate was added to the immune cells during FUS stimulation to mimic the tumoral background. Then, CD8 T cells or NK cells were co-cultured with H22 cancer cells or pEGFP-C1 plasmid-transfected Hepa1-6 cancer cells at a ratio of 1:1 in transwell devices (Corning Incorporated, NY, USA) with 3-µm diameter holes, which separated immune cells in the superstratum and cancer cells in the substratum for 48 h. Next, cells were observed using GFP fluorescence by the Zeiss observer3, crystal violet staining, and CCK-8 assay (Cat. No. ABS50003-500T, Univ, China) to assess cancer cell suppression and immune cell viability. Besides, the proportion of Ca^2+^ positive cells was monitored by FCM after incubation with the Fluo-4 AM dye.

### 2.9 Cytokine assay

The quantification of cytokine/chemokine in plasma was performed by Luminex xMAP technology using a magnetic Luminex assay (R&D Systems, Minneapolis, MN, USA), a Luminex® 200 Flow Cytometry System (Cat. No. x-200, Thermo Fisher Scientific, MA, USA), and Milliplex Analyst software (Version 5.1, Merck Millipore, MA, USA).

### 2.10 Preparation, characterization and therapeutic evaluation of STNDs@Ca^2+^

STNDs@Ca^2+^ were synthesized via ultrasonic emulsification in a fourth-step process. First, a lipid film was prepared by evaporating a chloroform solution containing DPPC, (FITC-)DOPE, Cholesterol, DSPE-PEG-HA, DSPE-mPEG2000, C9F19-COOH, and DPPA (purchased from Xi’an ruixi Biological Technology Co., Ltd) at the molar ratio of _1: 1: 2: 6: 1: 1: 1, followed by hydration to form a 1 mg/mL suspension. Second, 150 µL perfluoropentane (C_5_F_12_; Alfa, Zhengzhou, China) was added to 4 mL lipid suspension and sonicated to produce precursor STNDs@Ca^2+^ (pre-STNDs@Ca^2+^), which were size-refined by centrifugation and extruded to achieve 200∼300 nm. Third, calcium chloride solution (10 mmol/mL) were electrostatically complexed with pre-STNDs@Ca^2+^ to form primary STNDs@Ca^2+^ (pri-STNDs@Ca^2+^). Fourth, DC-Cholesterol·HCl (Ruixibio, Xi’an, China) was conjugated to pri-STNDs@Ca^2+^ to yield electroneutral STNDs@Ca^2+^, followed with purification by centrifugation at 6000 rpm and resuspended at ∼1×10¹² particles/mL for further use.

The size distribution and zeta potential of STNDs@Ca^2+^ were measured using the Malvern Nano Zetasizer (Malvern Instruments Ltd., Malvern, Worcestershire, U.K.), while the morphology was observed by laser confocal fluorescence microscopy (LCFM; LEXT OLS4000, Olympus, Tokyo, Japan) and cryo-transmission electron microscope (cryo-TEM; Krios G4 Cryo-TEM, Thermo Fisher Scientific, Massachusetts, USA).

To observe STNDs@Ca^2+^ biodistribution, following mice intravenously injected with 200 µL STNDs@Ca^2+^, visceral organs were subjected to FITC fluorescence imaging by IVIS LUMINA living imaging system (ILLIS; Caliper Life Sciences, California, USA) at 1-hour intervals over a 3-hour period, while sliced for imaging by the observer3 fluorescent microscopy.

To assay STNDs@Ca^2+^ toxicity, mice were intravenously injected with 200 μL STNDs@Ca^2+^ or saline once every two days for 5 times, and then blood was collected via retro-orbital puncture under isoflurane anesthesia (RWD, Shenzhen, China) and centrifuged at 3,000 × g for 15 minutes. Serum was isolated and analyzed using a biochemical autoanalyzer to quantify creatine kinase (CK), alanine aminotransferase (ALT), aspartate aminotransferase (AST), total bilirubin (TBIL), creatinine (CREA), uric acid (UA), and blood urea nitrogen (BUN). Additionally, visceral organs were harvested, fixed in paraffin, and subjected to HE and TUNEL staining.

One hour post-intravenous injection of STNDs@Ca^2+^, the experimental mice were subjected to FUS sti. spleen at 2.1 MPa to evaluate therapeutic efficacy of ultrasound-mediated spleen-targeted facilitation of exogenous Ca^2+^ influx activating splenic immunocompetence to suppress tumor proliferation. Detailed experimental procedures can be referred to in Section 2.2.

### 2.11 Statistical Analysis

Statistical analyses were conducted using GraphPad Software (version 8.0.1; GraphPad Software Inc., La Jolla, California, USA). Two-group comparisons were made using a two-tailed Student’s t-test assuming unequal variances. Regarding to multiple-group (>2 groups) comparisons, one-way analysis of variance (ANOVA) with Bonferroni’ multiple comparisons test was employed. Results were expressed as average ± standard error of mean (SEM) from at least three independent experiments. Differences were determined to be statistically significant at p < 0.05 (e.g., *, p<0.05; **, p<0.01; and ***, p<0.001).

## 3 Results

### 3.1 Antineoplastic efficacy and immunoregulation of SUS in orthotopic H22/Hepa1-6-HCC models

To elucidate the therapeutic efficacy and immunomodulatory properties of FUS sti. spleen, orthotopic H22/Hepa1-6-HCC mice were randomly allocated into three experimental cohorts: G1 and T1 group with normal mice classified as negative control (named normal), G2 group with orthotopic H22-HCC mice and T2 group with orthotopic Hepa1-6-HCC mice used as the positive control (named control), and G3 group with orthotopic H22-HCC mice and T3 group with orthotopic Hepa1-6-HCC mice respectively subjected to SUS (named FUS sti. spleen).

Regarding to G3 group, FUS sti. spleen started from day 2 post-H22 cell implantation into the liver, and performed once every two days for 20 days (Figure 1 D). Samples were collected on day 21 for follow-up experiments. In addition, another three groups of mice were subjected to the aforementioned procedure to calculate the survival rate for a period of 40 days. Anatomical images, tumor volume, and tumor weight showed that FUS sti. spleen significantly suppressed the proliferation of xenograft H22 carcinoma *in situ*, with a tumor suppression rate of up to ∼72% as compared with that in the control group (Figure 1 E-G; Supplementary Figure 1 A). More particularly, the volume of malignant ascites in the FUS sti. spleen group was remarkably lower than that in the control group, further suggesting the significant anticancer effect of FUS sti. spleen (Figure 1 H). Additionally, the liver weight (total weight of tumor and liver parenchyma) was significantly decreased in the G3 group compared to the G2 group, also indicating that SUS treatment significantly suppressed orthotopic H22-HCC tumor proliferation (Figure 1 I). The spleen weight showed a decreasing trend of splenomegaly in the G3 group as compared with the G2 group (Figure 1 J). The mice weight in the G3 group was lower than that in the G2 group, which also demonstrated that FUS sti. spleen significantly inhibited ascites production (Figure 1 K). More importantly, the survival curve showed that orthotopic H22-HCC mice subjected to therapeutic FUS sti. spleen had a longer survival time than those in the control group (Figure 1 L).

As regard to Hepa1-6-HCC treatment, T3 group was subjected to FUS sti. spleen starting from the second day post-cancer cell implantation, with treatments administered every other day for a duration of 30 days (Figure 1 M). Additionally, another three groups were subjected to the aforementioned procedure to calculate the survival rate for a period of 70 days. Anatomical images, tumor weight and tumor volume demonstrated significant orthotopic Hepa1-6-HCC tumor suppression attributed to therapeutic FUS sti. spleen, with a tumor inhibition rate of up to ∼83% as compared with that in the control group (Figure 1 N-P; Supplementary Figure 1 B). The liver weight of the T3 group was significantly lower than that of the T2 group, which further proved the significance of SUS in suppressing tumor proliferation (Figure 1 Q). In addition, spleen weight demonstrated that FUS sti. spleen significantly alleviated the symptoms of tumor-induced splenomegaly (Figure 1 R). The mice weight in the T3 group was lower than that of mice in the T2 group, which also demonstrated that FUS sti. spleen suppressed tumor proliferation (Figure 1 S). What’s more, Kaplan-Meier survival analysis indicated that SUS significantly extended the overall survival duration in orthotopic Hepa1-6-HCC tumor-bearing mice (Figure 1 T).

The FCM results showed that the orthotopic H22-HCC mice experienced a reduced proportion of crudely classified positive immune cells observably, such as CD4 T cells, Th2 cells, CD8 T cells, and DC1 in the spleen, CD4 T cells, Th1 cells, Th2 cells, NK cells, CD8 T cells, Mφ (Mφ2), and DC1 in the blood, and CD4 T cells, Th1 cells, CD8 T cells, Mφ1, and DC1 in the para-carcinoma tissues (Table 1; Supplementary Figure 2-4). Moreover, the proportions of traditional cognitive immunosuppressive cell populations, such as Th17 cells, Treg cells, and MDSCs (M-MDSCs and PMN-MDSCs) in both spleen and blood, and M-MDSCs in the H22-HCC para-carcinoma tissue were significantly increased (Table 1; Supplementary Figure 2-4). Regarding to the immune alteration in G3 group compared with the G2 group, FUS sti. spleen significantly activated host anti-cancer immunity. Concretely, the proportion of positive immune cells was significantly increased in the body, such as CD4 T cells, Th1 cells, Th2 cells, NK cells, CD8 T cells, Mφ2, and DC1 in the spleen, CD4 T cells, Th1 cells, Th2 cells, NK cells, CD8 T cells, Mφ (Mφ1 and Mφ2), DC1, and DC2 in the blood, CD4 T cells, Th1 cells, B cells, NK cells, CD8 T cells, Mφ1, and DC1 in the para-carcinoma tissue, and Th1 cells, NK cells, CD8 T cells, Mφ1, and DC1 in the tumor (Table 1; Supplementary Figure 2-5). Regarding the immunosuppressive cells, the proportions exhibited a significant reduction following SUS treatment, including Th17 cells, Treg cells and MDSCs (M-MDSCs and PMN-MDSCs) in the spleen and blood, Th17 cells and MDSCs (M-MDSCs and PMN-MDSCs) in the para-carcinoma tissue, and Th17 cells and PMN-MDSCs in the tumor (Table 1; Supplementary Figure 2-5). Additionally, the rate of Ki67 and cyclin-D1 positive cells in the spleen of the FUS sti. spleen group was higher than that in the control group, indicating that ultrasonic stimulation significantly promoted the proliferation of splenic cells, such as NK1.1 cells and CD8a positive cells (Figure 1 U). Moreover, the proportion of Ki67 and cyclin-D1 positive cells in the tumor of the FUS sti. spleen group was significantly lower than that in the control group, while NK1.1 cells and CD8a positive cells were significantly increased (Figure 1 V), which was in agreement with FCM results indicating that FUS sti. spleen facilitated splenic cells migration (i.e., NK cells and CD8 T cells) to the tumor to suppress cancer cell proliferation.

**Table 1.**
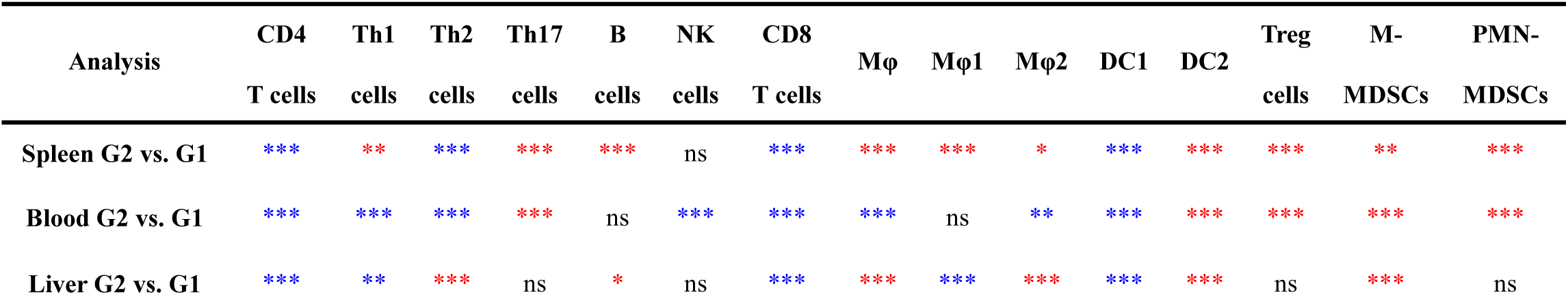
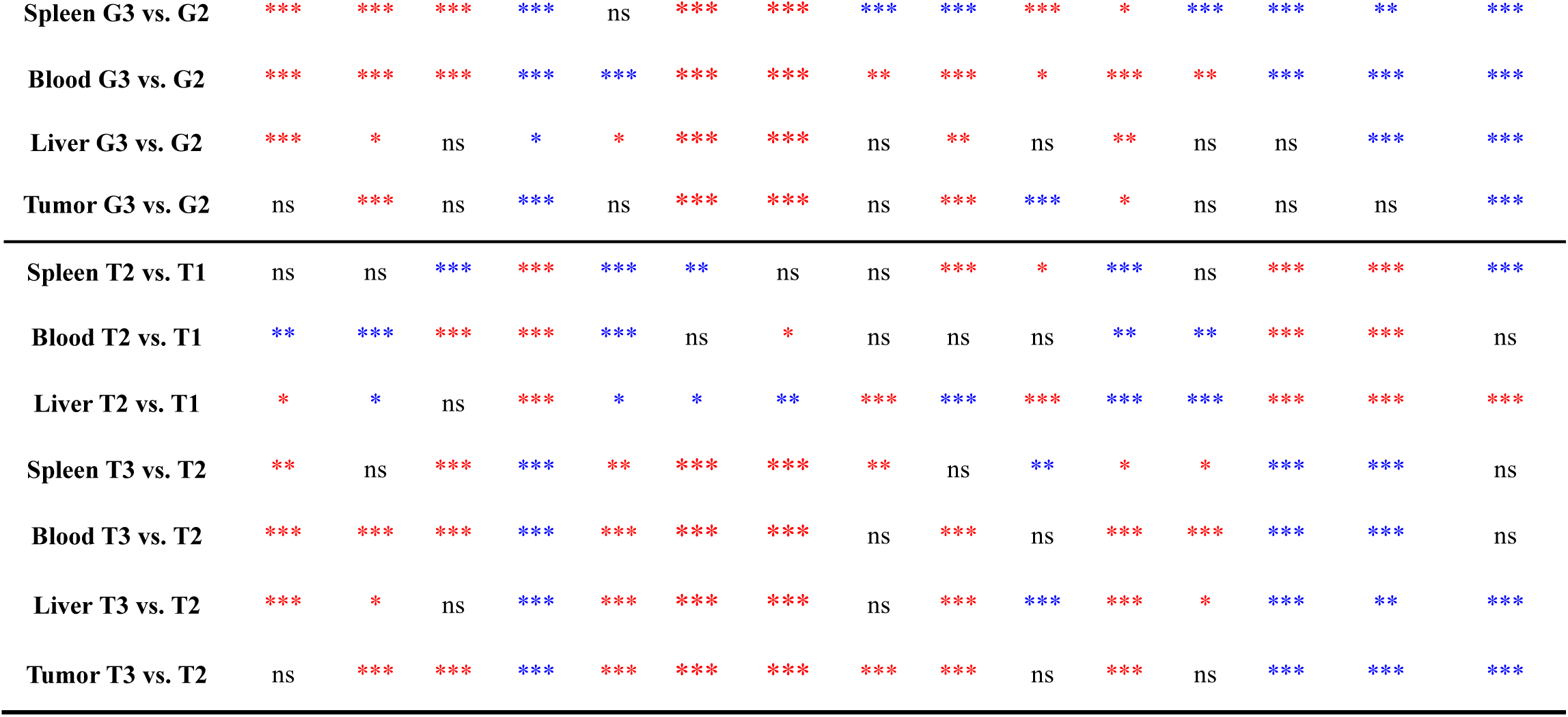
Statistical significance of FCM results of various immunocytes across the spleen, blood, tumor and para-carcinoma tissues from orthotopic H22/Hepa1-6-HCC mice subjected to FUS sti. spleen, non-treated murine tumor models, and normal mice. G1, normal mice; G2, control group of xenograft H22 HCC *in situ*; G3, FUS sti. spleen-treated orthotopic H22 HCC. T1, normal mice; T2, control group of xenograft Hepa1-6 HCC *in situ*; T3, FUS sti. spleen-treated orthotopic Hepa1-6 HCC. The red asterisk indicates a significant increase in the immunocyte proportions, while the blue asterisk indicates a significant decrease in the immunocyte proportions, and the ns represents no significant difference. The representative FCM plots and statistical diagram are shown in Supplementary Figure 2-9.

In orthotopic Hepa1-6-HCC mice, the proportions of immunosuppressive cells, including Th17 cells, Treg cells, and M-MDSCs, were consistently elevated in the spleen, blood, and peritumoral tissues, while the proportions of B cells and DC1 were significantly reduced (Table 1; Supplementary Figure 6-8). The proportions of other immune cell-types exhibited significant alterations in only 1-2 tissues, with Th2 cells and NK cells significantly reduced in the spleen, CD4+ T cells, Th1 cells, and DC2 markedly decreased in the peripheral blood, and Th1 cells, NK cells, CD8+ T cells, Mφ1, and DC2 significantly diminished in peritumoral tissues (Table 1; Supplementary Figure 6-8). Once orthotopic Hepa1-6-HCC mice subjected to FUS sti. spleen, the proportions of B cells, NK cells, CD8 T cells, and DC1 were significantly increased in the spleen, blood, peritumoral, and tumor tissues (Table 1; Supplementary Figure 6-9). Concurrently, CD4 T cells, Th1 cells, Th2 cells, Mφ1, and DC2 were markedly elevated in three tissue types (Table 1; Supplementary Figure 6-9). In contrast, the proportions of Th17 cells, Treg cells, and M-MDSCs were consistently and significantly reduced across all four tissues, followed by significant decreases in M-MDSCs and Mφ2 in two tissue types (Table 1; Supplementary Figure 6-9). Moreover, immunohistochemical analysis further demonstrated that FUS sti. spleen elicited a significant increase in the proportion of Ki67- and cyclin-D1-positive cells in spleen, indicating that ultrasonic stimulation robustly enhanced the proliferation of splenic immune cells, particularly NK1.1^+^ natural killer (NK) cells and CD8a^+^ cytotoxic T cells (Figure 1 W). Conversely, in tumor tissues, the proportions of Ki67- and cyclin-D1-positive cells were markedly reduced in the FUS sti. group, accompanied by a significant increase in NK1.1^+^ cells and CD8a^+^ T cells (Figure 1 X).

Collectively, these findings provided compelling evidence that FUS sti. spleen exerted profound immunomodulatory effects on splenic immunity, establishing its therapeutic potential as a novel immunotherapeutic strategy for malignant neoplasms. Particularly, SUS dramatically increased the proportions of NK cells, CD8 T cells, and DC1 in the spleen, blood, tumor, and para-carcinoma tissues consistently, followed by CD4 T cells, Th1, Th2, and Mφ1, whilst it reduced the proportions of Th17 cells, Treg cells and MDSCs. Concomitantly, the alterations in the immunocyte proportions across four tissues demonstrated that FUS sti. spleen facilitated the migration of splenic immunocytes, such as NK cells and CD8 T cells, to the tumor microenvironment. This immune cell redistribution ultimately contributes to the suppression of cancer cell proliferation.

### 3.2 scRNA-seq profiling splenic and intratumoral immune landscape responding to therapeutic FUS sti. spleen

Although FCM identified distinct lymphocyte heterogeneity across spleen, blood, tumor, and para-carcinoma tissues following FUS sti. spleen, this methodology presented limitations in comprehensively characterizing the dynamic evolution of cellular subpopulations within identical lineages. Furthermore, it failed to elucidate the inherent immunogenic heterogeneity exhibited by diverse cellular subtypes. Thereafter, scRNA-seq was performed to delineate the splenic and intratumoral multifaceted landscapes from G2 and G3 groups. Based on the comprehensive FCM data (Table 1), which demonstrated that CD8+ T cells and NK cells exhibited the most pronounced changes in proportion across all examined tissues in response to SUS, coupled with extensive literature highlighting the pivotal role of T cells and NK cells in killing cancer cells and antitumor immunity, we focus herein on presenting the scRNA analysis of T_NK cells, while detailed findings regarding other cell types are provided in Supplementary File 1.

#### 3.2.1 SUS altered the proportion and physiological function of splenic T_NK cells

Unsupervised clustering of all T_NK cells was performed using the spectral clustering method to reveal the intrinsic structure and potential functional subtypes of the overall T_NK cell populations [32]. A total of 19 stable clusters were found according to the differential expression of unique signature genes, including 8 clusters for CD4 T cells (C01-C08), 6 clusters for CD8 T cells (C09-C14), 2 clusters for proliferative T cells (C15-C16), and 3 clusters for NK cells (C17-C19; Figure 2 A-B). According to the profiling of top 10 uniquely expressed genes across T_NK subclusters (Supplementary Figure 10 A), the “inhibitory” marker Ctla4 was highly expressed in the CD4 T cell subtypes C01-C02 and C06, which was a key modifier of CD4 T cells to protect against maladaptive cytotoxicity [33], while Foxp3 was specially expressed in C06. Lef1 was significantly expressed in C03-C05 and C08 subclusters, which inhibited the increase of Treg suppressive function after the activation and promoted tumor-infiltrating CD8 T cell persistence [34, 35]. Additionally, the Trdc gene as γδT marker was highly expressed in C07, which had extraordinary antitumor potential and prospects [36, 37]. The “naive” marker genes such as Lef1 and Ccr7 were specially expressed in the C9-C11 of the CD8 T clusters [38]. High expression of Ccl5 and Ctla2a was found in the C12-C14 subclusters, which were commonly associated with T cell effector functions [39, 40]. Kif18b and Tal1 genes were significantly upregulated in C15 and C16, respectively, which were commonly related to T cell proliferation [41–44]. Notably, specific expression of cytotoxic markers, such as Thy1, Gzmb, and Klri2, was found in the three NK cell subtypes C17-C19 [45, 46], commonly related to the regulation of NK cell-mediated cytotoxicity [47]. Particularly, the cell proportions of C04, C11-C12, C14, and C16-C18 subclusters were increased in the FUS sti. spleen group compared to the control group, while that of the others were decreased (Figure 2 C). The modulation in cellular proportions within these subclusters, characterized by either a significant reduction or increment, demonstrated their responsiveness to SUS, suggesting a dynamic and potentially regulated interaction between the FUS stimulus and cellular behavior. It is noteworthy that, in response to FUS sti. spleen, the cell proportions of C01-C02 and C06 as “inhibitory” immunocytes were decreased, while that of “cytotoxic” C12, C14, and C17-C18 was increased (Figure 2 C), suggesting the significant alteration of splenic immunity by FUS. According to tumorigenic progression, and ultrasound-mediated dynamic splenic immunomodulation, we postulate with substantial biological plausibility that ultrasound preferentially modulates splenic immunity through two plausible mechanisms: (1) with great probability, SUS attenuated tumor-driven immunosuppressive polarization, accompanied by activation of splenic immunocyte subsets, or alternatively (2) phenotypic reprogramming of immunosuppressive niches towards immunocompetent states. Such hypothesis may represent a distinct yet promising avenue for subsequent exploration, but it extends beyond the primary research objectives delineated in this study.

**Figure 2.**
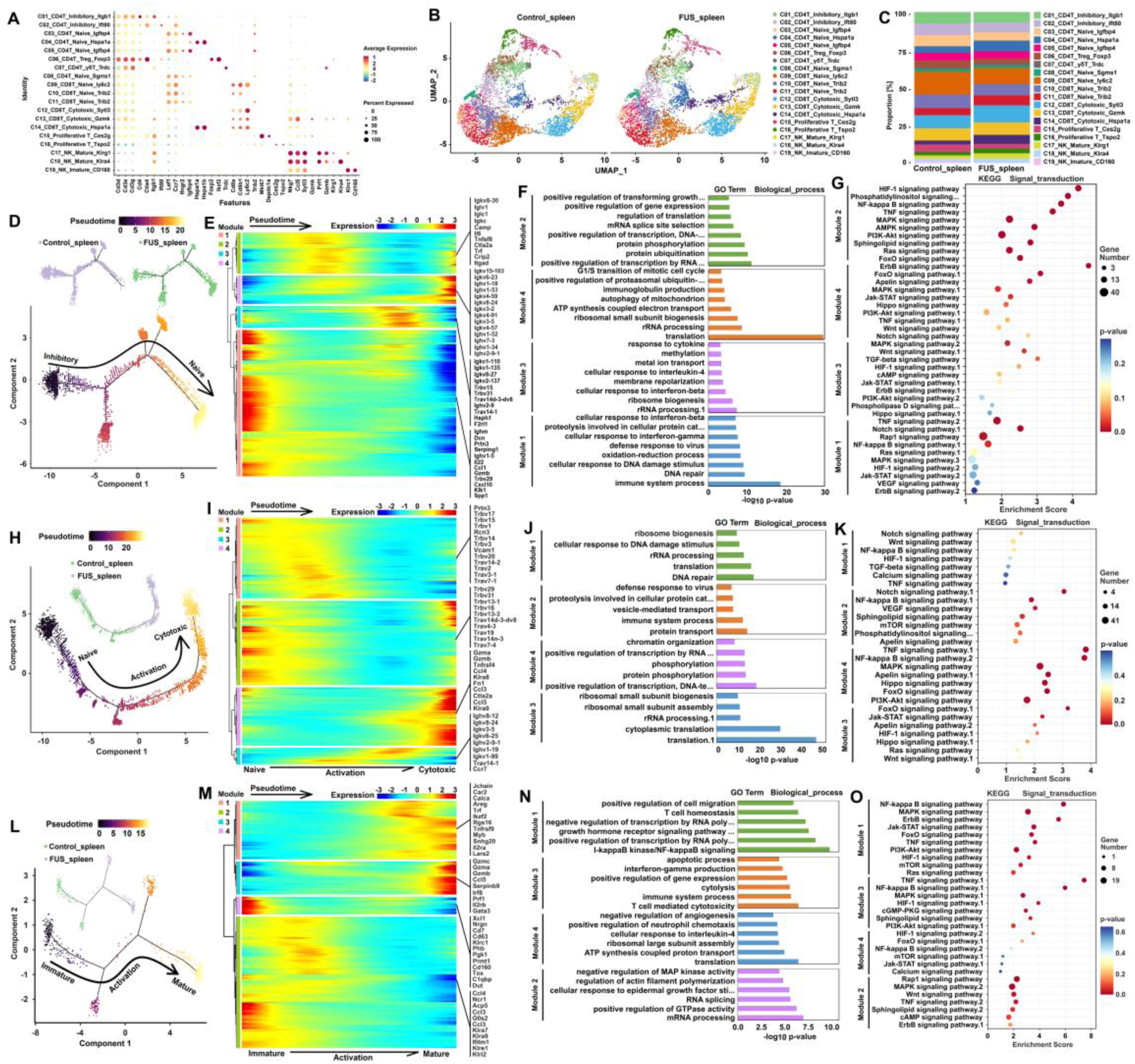
FUS sti. spleen significantly improved the proportion and physiological function of splenic T_NK cells. A, dot plots of canonical marker genes applied for T_NK cell subclusters identification. B, UMAP projection of T_NK cells from both the control and FUS sti. spleen group. C, comparison of T_NK cell proportion between the control group and FUS sti. spleen group. D, H, and L, pseudotime-ordered analysis of CD4 T cells, CD8 T cells, and NK cells, respectively, from the control group and FUS sti. spleen group. E, I, and M, pseudotime heatmap showing the dynamic changes in gene expression of CD4 T cell, CD8 T cell, and NK cell subclusters, respectively, based on their common kinetics through pseudotime using Monocle 2. Genes (rows) were clustered into 4 modules, and cells (columns) were ordered according to pseudotime. F, J, and N, distinct GO terms of biological process and their *p* value associated with each module in CD4 T cells, CD8 T cells, and NK cells respectively. G, K, and O, KEGG signaling pathway enrichment of signal transduction in CD4 T cells, CD8 T cells, and NK cells respectively.

Pseudotime trajectory analysis was then performed to elucidate the dynamic cell transition of T_NK cells responding to FUS sti. spleen or not. The cell trajectory mainly started from C01, C02, and C06 subtypes and ended with C03, C05, and C08, suggesting a suppression of the “naive” to “inhibitory” transformation during the tumor progression attributed to FUS sti. spleen (Figure 2 D; Supplementary Figure 10 B). The profile of gene expression dynamics identified 4 modules across the pseudotime trajectory (Figure 2 E). Along with the pseudo-chronological timeline, the genes of Camp, Il6, Trf, Crip2, and Itgad in the module 2 related to T cell proliferation, activation, and motility were significantly upregulated as a consequence of FUS stimulation. In addition, module 4 gradually expressed high levels of Igkv and Ighv family members that were enriched in immunoglobulin production and immune process. SUS significantly promoted gene expression associated to the GO terms of ribosome biogenesis, transcription and translation, and protein modification (Figure 2 F). KEGG enrichment analysis further revealed the upregulation of HIF-1, TNF, NFκB, phosphatidylinositol, MAPK, and ErbB signaling pathways, which regulated the antitumor immune response (Figure 2 G) [48–50].

The cellular state evolved from naive to cytotoxic state according to the CD8 T cell subtypes and proportion mapped on the pseudotime trajectory (Figure 2 H; Supplementary Figure 10 C). The significantly changed genes among CD8 T cell subclusters in the pseudotime trajectory were examined and arranged into 4 modules with distinct GO functions and KEGG enrichment according to their pseudotemporal expression patterns (Figure 2 I-K). GO enrichment analysis of gene sets showed an enhancement in cellular transcription and translation along the pseudotime axis that responded to FUS stimulation (Figure 2 J). The Notch signaling pathway was significantly enriched especially in initial period of modules 1 and 2, but was significantly inhibited at the posterior timeline (Figure 2 K), suggesting the transformation of CD8 T cell naivety, T cell dysfunction, and inert immune response towards effector function [51]. Moreover, the temporal expression profile of TRAV and TRBV subfamily genes exhibited a biphasic pattern characterized by initial downregulation followed by subsequent upregulation (Figure 2 I), suggesting the oscillatory diversity and clonality repertoire of CD8 T cells, which potentially reflects the immunological adaptation and functional reprogramming of T cells under FUS modulation [52]. Particularly, TNF, NFκB, and FoxO signaling pathways were enriched in modules 3 and 4 (Figure 2 K), which exerted cytotoxic function on cancer cells [53–55]. Meanwhile, it is accompanied by high expression of the cytotoxic and proliferation markers such as Klra8, Gzma, Gzmb, Gzmk, Sfn, and Trbc, Trgv, Igkv, Ighv, and Klra family members along the trajectory on modules 3 and 4, demonstrating the proliferation and activation of splenic CD8 T cells under FUS stimulation (Figure 2 I).

The pseudotime trajectory analysis of NK cells revealed that SUS promoted immature C19 subtype of NK cells evolving to the mature C17 and C18 but may also promote the proliferation of NK cells (Figure 2 L; Supplementary Figure 10 D). Predicted gene expression was plotted in pseudotime to track changes across different NK cell states, revealing 4 modules with distinct GO functions and KEGG enrichment (Figure 2 M-O). Particularly, the module 3 was significantly different in high expression of the cytotoxic genes Gzmb, Gzmc, Gzma, Irf8, and Prf1 (Figure 2 M) along the trajectory that was enriched in GO terms related to regulation of NK cell activation and immune process (Figure 2 N). The KEGG enrichment analysis also revealed that the TNF, NFκB, and MAPK signaling pathways were enriched in modules 1 and 3, indicating a response to FUS stimulation (Figure 2 O). Collectively, these results fully proved that FUS sti. spleen activated T_NK cell immunocompetence and transformed these immunocyte immunosuppression to suppress HCC proliferation.

#### 3.2.2 SUS transformed intratumoral T_NK cell landscape suppressing cancer cell deterioration

A deep characterization of the tumor-infiltrating T_NK cell landscape would provide better understanding of the action mechanism of FUS sti. spleen in the eradication of cancer. The therapeutic effect on tumor was at least in part attributed to the cytotoxic T_NK cells in the TME, but the cytotoxic activity of these cells could also be ineffective primarily by the suppression of Tregs, MDSCs or the sustained expression of inhibitory receptors that lead to a T_NK cell dysfunction state [56–59]. Therefore, the intratumoral T_NK population was analyzed with 14 distinct subclusters (Figure 3 A-B), including four distinct subtypes of CD4^+^ T cells (C01-C04), five main subpopulations of CD8^+^ T cells (C05-C09), three subclusters of CD4-CD8- double negative T cells (DNT; C10-C12), and two subclusters of NK cells (C13-C14). As depicted in heatmap plotting the top 10 specially expressed genes across intratumoral T_NK cell subclusters (Supplementary Figure 11 A), the “inhibitory” marker genes Ctla4 and Tigit were highly expressed in the CD4 T cell subtypes of C01, C02, and C04, and Foxp3 was specially expressed in the C04 [33, 60]. Remarkable expression of the cytotoxic or effector marker genes Gzmk, Nkg7, and Ccl5 was found in the CD8 T cell subclusters of C05-C09 [39, 40], but the “inhibitory” markers Ctla4 and Tigit were upregulated in C08-C09 [33]. The “naive” marker genes Lef1 and Ccr7 were expressed in the DNT subcluster of C10 [38]. Additionally, the IL-17 family members Il17a and Il17f, usually bond to receptors in TME linking to regulate tumor growth and metastasis [61], were highly expressed in the C11 subcluster. The Trdc gene was highly expressed in the C12 DNT subcluster, which indicated an extraordinary antitumor potential and prospects [37]. Particularly, the NK cell subtypes C13-C14 were characterized by the specific expression of Gzma, Gzmb, Nkg7, and Ccl5 that were usually related to the regulation of NK cell-mediated cytotoxicity and effector function [45–47], but the inhibitory KIR family members, which abrogated NK cell activation by binding to their major histocompatibility complex (MHC) class I ligands [62], were highly expressed in both tumoral C13-C14 populations. Notably, the cell proportion of C04, C06, C07, C10, and C12 was increased, while that of the others were decreased (Figure 3 C). Although the cell proportion of only C06 and C07 cytotoxic CD8 T cells was significantly increased while other T_NK cell subpopulations decreased or showed no change, it should be noted that the proportion of inhibitory T cells and infaust cell populations in FUS sti. spleen group was much less than that in the control group. Combined, the TME in the control group containing CD4 T cells, CD8 T cells, and NK cells displayed an immunosuppressive state, but FUS sti. spleen attenuated tumor-driven immunosuppressive polarization of T_NK cells, and particularly increased the proportion of tumor-infiltrated cytotoxic CD8 T cells.

**Figure 3.**
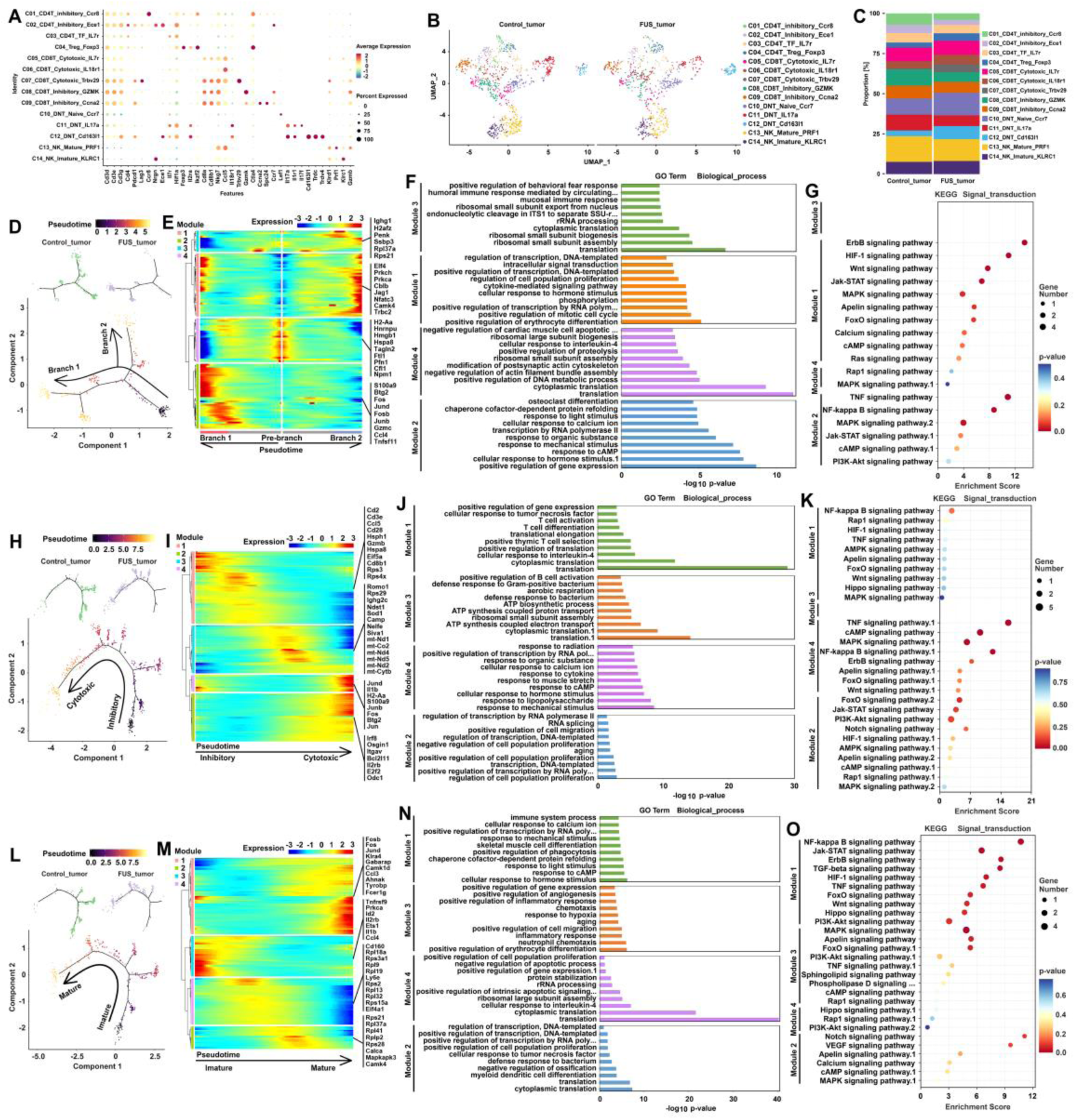
SUS significantly modified T_NK cells in TME from a negative to a positive tumor suppressive state. A, dot plots of signature markers applied for intratumoral T_NK cell subclusters identification. B, UMAP projection of intratumoral T_NK cells from the control group and FUS sti. spleen group. C, cell proportion of intratumoral T_NK cell subclusters between the control group and FUS sti. spleen group. D, H, and L, pseudotime-ordered analysis of CD4 T cells, CD8 T cells, and NK cells, respectively, from the control group and FUS sti. spleen group. E, I, and M, pseudotime heatmap showing the dynamic changes in gene expression of CD4 T cell, CD8 T cell, and NK cell subclusters, respectively, based on their common kinetics through pseudotime using Monocle 2. Genes (rows) were clustered into 4 modules, and cells (columns) were ordered according to pseudotime. F, J, and N, distinct GO terms of biological process and their *p* values associated with each module in CD4 T cells, CD8 T cells, and NK cells, respectively. G, K, and O, KEGG signaling pathway enrichment of signal transduction in CD4 T cells, CD8 T cells, and NK cells, respectively.

When CD4 T cells were delineated in pseudotime, the C01-04 subclusters of CD4 T cells were categorized into 2 fates (branches 1 and 2) along the trajectory (Figure 3 D; Supplementary Figure 11 B). The mechanical stimulus-related genes Btg2, Fos, Jund, Fosb, and Junb were significantly upregulated in the module 2 of branch 1 as dissected in gene expression patterns (4 modules) involved in the continuum transition (Figure 3 E). In particular, the genes in the module 1 related to immune cell proliferation and activation, such as Elf4, Cblb, Jag1, and Nfatc3 (Figure 3 E), were gradually upregulated along with the trajectory differentiation process. Thus, the intratumoral CD4 T cells from FUS sti. spleen group were enriched in GO terms related to positive regulation of gene expression and erythrocyte differentiation (Figure 3 F), which were correlated with the modulation of TNF, and NFκB, ErbB, and HIF-1 signaling pathways (Figure 3 G).

Next, the dynamic state transition of CD8 T cells was explored by investigating the pseudotime trajectory using Monocle 2 (Figure 3 H; Supplementary Figure 11 C). This analysis showed that the “inhibitory” CD8 T cells (C08-C09 subclusters) were at the beginning of the trajectory, whereas “cytotoxic” CD8 T cells were in a terminal state (Supplementary Figure 11 C). Four modules of differentially expressed genes along the CD8 T cell trajectory were identified (Figure 3 I). The module 3 showed high expression of mt gene family members, involved in the synthesis of ATP in the mitochondria, along the trajectory, and the module 2 showed the upregulation of Irf8, Osgin1, Bcl2l11, and Odc1 that were related to T cell population proliferation, activation, and exhaustion (Figure 3 I-J). Moreover, Jund, Fosb, Fos, Btg2, and Jun genes involved in the GO term of response to mechanical stimulus were enriched in the module 4 (Figure 3 I-J), which mainly involves the intracellular regulation of TNF, cAMP, MAPK, and NFκB signaling pathways by ultrasonic stimulation (Figure 3 K). Accordingly, these observations suggested that FUS sti. spleen attenuated tumor-driven immunosuppressive polarization of intratumoral CD8 T cell subsets, accompanied by immune activation. Moreover, the proportion of cytotoxic cells was increased after therapy, and that of the inhibitory ones was decreased (Figure 3 C).

The trajectory analysis of the intratumoral NK cells based on the Monocle 2 algorithm was also performed to assess NK cell transition in H22-HCC lesions (Figure 3 L). The NK cells were categorized into 4 states along the trajectory, where the cell proportion of state 3-4 indicated that the “mature” phenotype was significantly increased (Supplementary Figure 11 D). The genes (such as Fosb, Fos, Jund, Klra4, Camk1d, Ccl3, Tyrobp, and Fcer1g) related to cellular response to mechanical stimulus, NK cell proliferation and activation, and immune process were significantly enriched, whereas the genes related to intracellular transcription and translation were significantly reduced as dissection of dynamic gene patterns involved in the NK cell state transition (Figure 3 M-N). KEGG enrichment analysis further revealed that NFκB, MAPK, Jak-STAT, and ErbB signaling pathways were significantly upregulated during the process of NK cell maturation (Figure 3 O).

Notably, CNV analysis revealed that the epithelial cells exhibited the highest CNV level among the main cell populations, which was higher in the control group than in the FUS sti. spleen group (Figure 4 A). This suggested that the deterioration of cancer cells was effectively curbed after the orthotopic H22-HCC mice were subjected to FUS sti. spleen. Unsupervised clustering with t-SNE revealed 7 subclusters of epithelial cells, which were annotated by known or putative markers (Figure 4 B-C). All subgroups except for the C6 subcluster abnormally expressed oncogenic genes (including Large1, mt-Cytb, Calu, Aplp2, Adgrf5, and Plec; Supplementary Figure 11 E) [63–68]. In addition, all epithelial cell subclusters consistently overexpressed RPS and PRL gene family members that enhanced ribosomal protein content and ribogenesis contributing to tumor onset and progression (Supplementary Figure 11 E) [69, 70]. The cell proportion in C1 and C3-C5 increased following therapeutic FUS sti. spleen, but was reduced in C2 and C6-C7 (Figure 4 D). However, it is far more crucial that CNV levels of C1-C7 were reduced after FUS sti. spleen as compared to the control (Figure 4 E), suggesting that FUS sti. spleen restricted cancer’s malignant evolution.

**Figure 4.**
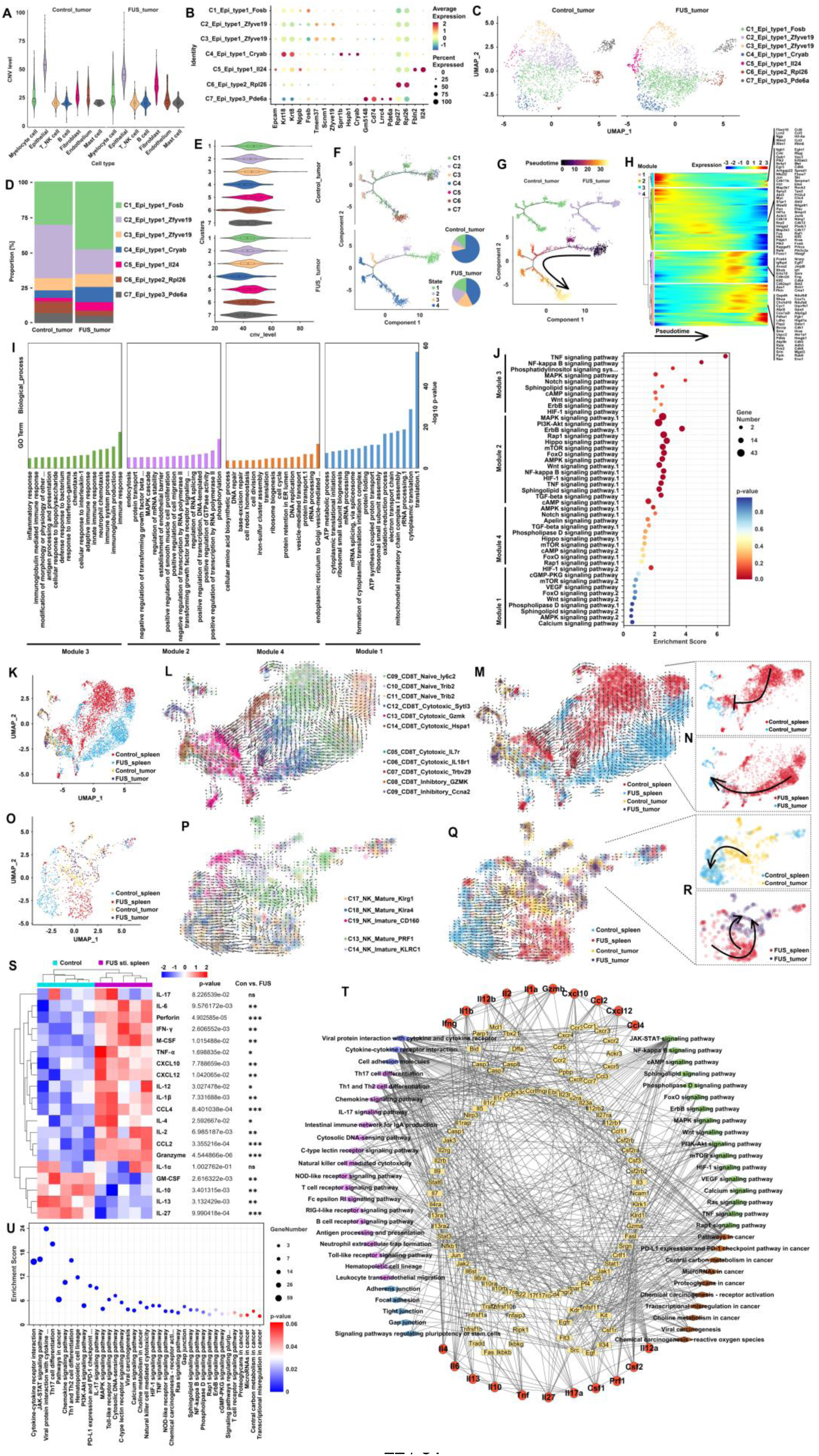
Characterization of epithelial cell landscape to evaluate tumor progression, pseudotime trajectory analysis of CD8 T cell and NK cell trafficking between spleen and tumor, and cytokine profiling in plasma. A, violin plot showing CNV level of distinct cell compositions in TME between the G2 and G3 groups. B, dot plot of various expression profiles of signature genes across epithelial cell subtypes. C, UMAP plot of epithelial cells stratified into seven distinct subclusters. D, the proportion of epithelial cell subclusters in the G2 group and G3 group. E, violin plot of CNV level across epithelial cell subclusters between the G2 group and G3 group. F, pseudotime identifying 4 states of epithelial cell subclusters along the trajectory, and their cell proportion analysis. G, pseudotime-ordered analysis of epithelial cells from the G2 group and G3 group. H, pseudotime heatmap showing the dynamic gene expression in the epithelial cell subclusters based on their common kinetics through pseudotime using Monocle 2. Genes (rows) were clustered into 4 modules, and cells (columns) were ordered according to pseudotime. I, distinct GO terms of biological process and their *p* values associated with each module in epithelial cells. J, KEGG signaling pathway enrichment of signal transduction in epithelial cells. K, UMAP projection of CD8 T cells from both spleen and tumor of both G2 and G3 groups. L-M, scVelo plot of CD8 T cell subclusters from both spleen and tumor of both G2 and G3 groups. N, scVelo plot of CD8 T cell subtypes in the G3 group compared to G2 group, which demonstrating CD8 T cell transition from spleen to tumor in response to FUS sti. spleen. O, UMAP projection of NK cells in from both spleen and tumor of both G2 and G3 groups. P-Q, scVelo plot of NK cell subclusters from both spleen and tumor from both G2 and G3 groups. R, scVelo plot of NK cell subclusters in the G2 group or G3 group, which demonstrating NK cell transition from spleen to tumor in response to SUS. S, heatmap of cytokine levels in plasma from orthotopic H22-HCC mice with or without FUS sti. spleen treatment. T, protein-protein network analysis of cytokines and target genes based on STRING database. U, KEGG pathway enrichment analysis of the target genes of cytokines.

Next, Monocle 2 pseudotime analysis was used to explore the dynamics of malignant evolution across epithelial cell subclusters (Figure 4 F-G). This analysis delineated 4 distinct cellular states, revealing a progressive transition from clusters C1, C2, C4, and C5 through intermediate C3 and C7, culminating in terminal state C6, which demonstrated a significant reduction in malignant cell proliferation and CNV burden in response to therapeutic SUS. The gene expression dynamics were categorized as 4 main pseudotime-dependent gene modules (Figure 4 H), which proved that FUS sti. spleen intervened in the tumor malignant trajectories. Module 2 consisted of various proto-oncogenes (including Crkl, Elk4, Myc, Fosb, Fos, Jun, Met, Ski, Akt3, and Raf1) that were upregulated in the early stage but downregulated in the end-stage of the trajectory (Figure 4 H). Besides, genes in modules 1 and 4, annotated to promote metabolism (including oxidative phosphorylation, glycolysis/gluconeogenesis, and glutathione metabolism), cell proliferation, and tumor metastasis, were upregulated at the intermediate-stage but displayed significant downregulation at the end (Figure 4 H-I), which further suggested FUS sti. spleen arrested and blocked both the level of metabolic reprogramming and the ability of cell deterioration. KEGG pathway enrichment analysis of four gene modules indicated suppression of TNF, MAPK, NFκB, PI3K-Akt, AMPK, and HIF-1 signaling pathways in epithelial cells after immunomodulation by therapeutic FUS sti. spleen (Figure 4 J).

In this work, scRNA-seq data from CD8 T cell or NK cell subpopulations in the spleen and tumors were pooled for a combined pseudotime trajectory analysis to further demonstrate the migration of splenic immune cells to tumors in response to FUS stimulation. During the analysis, CD8 T or NK cell subtypes in both spleen and tumor from the FUS sti. spleen group and control group were mapped separately, distinguishing their tissue origins (Figure 4 K, O). The scVelo plot of CD8 T cell subtypes, particularly in FUS sti. spleen group, clearly showed that the splenic “naive” C9-C11 subclusters transited to “cytotoxic” C12, C13, and then C14 subcluster, ending in tumor along the trajectory (Figure 4 L-M). Importantly, the transition process of CD8 T cells from spleen to tumor was significantly interrupted in the control group, in particular, the transition from C13 subcluster to C14 subcluster was blocked (Figure 4 N). In contrast, FUS sti. spleen facilitated the migration of “cytotoxic” CD8 T cells from spleen to tumor (Figure 4 M-N). In particular, the splenic C14 subcluster of “cytotoxic” CD8 T cells migrating to tumor showed enrichment of TCR clonotypes associated with anti-tumor effects (Supplementary File 1). Moreover, the scVelo plot of NK cell subtypes revealed that in the control group, mainly the “immature” C19 subcluster transited to the tumoral “mature” C13 subcluster (Figure 4 O-P). However, the scVelo plot of NK cell subtypes in the control group alone showed that NK cell evolution was directed from the tumor to the spleen, indicating immunosuppression (Figure 4 Q-R). In contrast, in the FUS sti. spleen group, the C17 and C18 subclusters of “mature” NK cells transited from spleen to tumor (Figure 4 Q-R). Overall, ultrasound activated splenic CD8 T cells and NK cells, and promoted their migration to tumor lesions.

In addition to cellular immunity, various cytokines are involved in both innate and adaptive immune responses to suppress tumor proliferation [71]. Herein, Luminex xMAP technology was used to selectively assess the levels of the following cytokines in the peripheral blood: GM-CSF, TNF-α, IL-12, CCL2, IL-1β, IL-2, IL-4, IL-6, IL-10, IL-13, IL-17, IFN-γ, CXCL10, M-CSF, IL-1α, CCL4, CXCL12, IL-27, perforin, and granzyme. The upregulated levels of TNF-α, IFN-γ, perforin, and granzyme further demonstrated the activation of CD8 T cells and NK cells in response to FUS sti. spleen (Figure 4 S) [72–74]. Besides, GM-CSF was generally correlated to tumor progression of glioblastoma, small cell carcinoma, skin carcinoma, meningiomas, colon cancer, lung cancer, and particularly HCC, by the regulation of the tumor microenvironment involving Mφ and MDSCs, and the promotion of epithelial to mesenchymal transition, angiogenesis, and expression of immune checkpoint molecules [75–78], which was significantly reduced after FUS sti. spleen (Figure 4 S). M-CSF level was increased after the treatment of orthotopic HCC mice with FUS sti. spleen (Figure 4 S), which regulated the survival, proliferation, and differentiation of the monocyte-macrophage lineage from progenitors to mature cells and activated several important functions of mature tissular Mφ [79, 80]. Additionally, ILs displayed pleiotropic roles in the pathogenesis and resolution of various tumors. The proinflammatory cytokine IL-1β exerted contradictory effects on primary tumors, but anti-cancer treatments promote IL-1β secretion [81, 82], which resembles the anticancer effect of FUS sti. spleen. IL-2 is a pleiotropic cytokine required for both effector lymphocyte proliferation/differentiation and regulatory T cell expansion/survival [83], which was also highly secreted as a consequence of spleen responding to FUS stimulation (Figure 4 S). Additionally, IL-4, IL-6, and IL-12 positively improved survival and effector functions of T_NK lymphocytes [71, 84, 85], but IL-10 and IL-27 are involved in immune tolerance and immunosuppression, such as promotion of Foxp3^+^ Treg cell formation, maintenance, and growth [86, 87]. IL-13 was most of the time considered as a cancer immunotherapeutic target by boosting type-1-associated anti-tumor defense [88]. In this work, the concentration of IL-4, IL-6, and IL-12 was increased but that of IL-10, IL-27, and IL-13 was decreased when the tumor mice were subjected to FUS sti. spleen (Figure 4 S). However, beyond that, the levels of IL-1α and IL-17 were not significantly different between the FUS sti. Spleen group and the control group (Figure 4 S). The increased levels of the pro-inflammatory chemokines CCL2, CCL4, and CXCL10 (Figure 4 S), promoted the migration of immune cells, especially effector T cells transiting into TME [89–91], which corroborates the aforementioned pseudotime trajectory analysis revealing the migration of CD8 T cells from the spleen to tumor. The upregulated CXCL12 (Figure 4 S) also promoted the migration and activation of most leukocytes, and usually synergized with CXCL8 but also with CXCL9, CXCL10, CXCL11, and multiple CC chemokines to attract B cells, T cells, DCs, monocytes, and other leukocytes [92–94]. Moreover, the protein-protein network analysis of cytokines and target genes based on STRING database, and KEGG pathway enrichment analysis demonstrated that SUS facilitated cytokines secretion against tumor via several signaling pathways (such as JAK-STAT, chemokine, and other signaling pathways), in particular, unequivocally regulating the cytotoxic potential of NK cells and CD8^+^ T cells through the cytokine-cytokine receptor interaction pathway (Figure 4 T-U).

### 3.3 SUS facilitated calcium influx into splenic immunocytes to revitalize anticancer immunocompetence

Accordingly, FCM results and scRNA analysis elucidating the dynamic cell proportions and transitions (involving T_NK cells, myeloid cells and B cells) demonstrated that FUS sti. spleen effectively mitigated tumor-induced immunosuppressive polarization of the spleen, concurrently activating splenic immunocyte subsets and facilitating their migration to suppress tumor proliferation (Table 1; Figure 2-4; Supplementary File 1). Unequivocally, ultrasound-mediated immunomodulation exerts non-specific effects on splenic cellular types or subtypes. In subsequent experimental investigations, we strategically focused on characterizing CD8 T lymphocytes and NK cells, as they exhibited the most substantial immunophenotypic alterations and usually play a paramount role in mediating tumoricidal activity through direct cellular cytotoxicity.

Following the comprehensive scRNA profiling, we systematically performed FCM using the signature marker CCL5 and CCR7 to quantify “naive” or “inhibitory” and “cytotoxic” CD8 T cell subtypes (Figure 2 A), and CCL5 plus CD159a (Klrc1) to profile “immature” and “mature” NK cell subtypes (Figure 3 A) in G1-G3 groups. Additionally, the classical markers CD137 (4-1BB) and IFN-γ were used to distinguish the “cytotoxic” CD8 T cells from those that were not, and perforin was used to distinguish “cytotoxic” NK cells from those that were not. The FCM results further proved that FUS sti. spleen significantly increased the proportion of “effector” CD8 T cells and NK cells, and promoted their transition from the immunosuppressive state to the activated state (Figure 5 A; Supplementary Figure 12). Additionally, GSVA results of the scRNA-seq data further confirmed the upregulation of TNF, NFκB, MAPK, HIF-1, PI3K-Akt, AMPK, ErbB, cAMP, Rap1, Jak-STAT, Calcium, and other signaling pathways across most CD8 T cell subtypes and NK cell subsets following FUS exposure (Figure 5 B-C), exhibiting remarkable concordance with the above pseudotime trajectory analysis (Figure 2 H-O). Such findings were further confirmed by the GSEA results of bRNA-seq of splenic CD45^+^ CD3^+^ CD8a^+^ cells (CD8 T cells) and CD45^+^ CD3^-^ NK1.1^+^ cells (NK cells) sorted from both G2 and G3 groups, showing significant upregulation of TNF, NFκB, MAPK, Calcium, and other signaling pathways in both cell types as a consequence of the ultrasound stimulation (Figure 5 D-G). Subsequently, based on integrated analysis and drawn upon well-established biophysical and mechanistic studies of cellular modulation by ultrasound, radiation, electrode, electromagnetism, and other physical means [95–97], we propose a plausible molecular cascade wherein FUS facilitated calcium influx subsequently triggering activation of the above mentioned signaling pathways (Figure 5 H), which synergistically orchestrate the observed immune activation phenotype.

**Figure 5.**
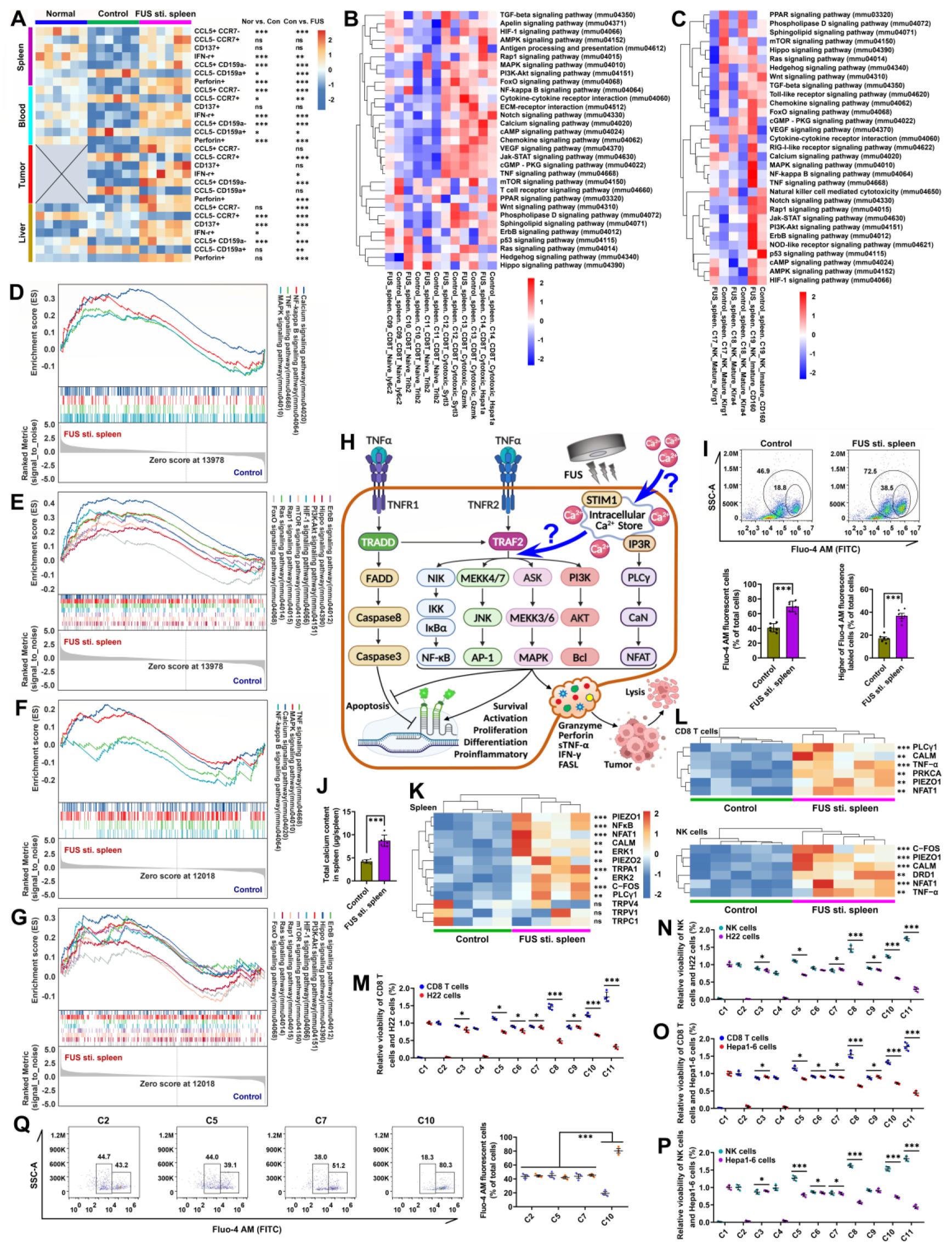
FCM analysis of activated CD8 T cells and NK cells in orthotopic H22-HCC mice subjected to FUS sti. spleen, and differential analysis of KEGG signaling pathways, as well as the results of Ca^2+^ enhanced cancer cell suppression *in vitro*. A, heatmap showing FCM results of “naive”, “inhibitory” or “cytotoxic” CD8 T cell subtypes in G1-G3 groups based on the signature genes of CCL5 and CCR7 displayed in Figure 2 A and 3 A, and “immature” or “mature” NK cell subtypes according to the signature genes of CCL5 and CD159a (Klrc1). Additionally, the classical markers CD137 (4-1BB) and IFN-γ were used to distinguish the “cytotoxic” CD8 T cells or not, and perforin was used to distinguish “cytotoxic” NK cells or not. The corresponding flow charts and statistical analysis are shown in Supplementary Figure 12. B-C, GSVA analysis of partial KEGG pathways enrichment analysis in splenic CD8 T cell subtypes and NK cell subtypes (as clustered in Figure 2 A-B), respectively. D-E, GSEA analysis of partial KEGG pathways enrichment analysis in splenic CD8 T cells. F-G, GSEA analysis of partial KEGG pathways enrichment analysis in splenic NK cells. H, schematic diagram of FUS enhancing calcium influx and regulating TNF, NFκB, MAPK, and other signaling pathways. I, flow charts of Fluo-4 labeled splenic cells from orthotopic H22-HCC mice with or without FUS sti. spleen treatment, and the corresponding quantification. J, assay of calcium content in the whole spleen. K-L, heatmaps showing qRT-PCR results in the spleen, splenic CD8 T cells, and splenic NK cells of orthotopic H22-HCC mice with or without FUS sti. spleen treatment. M-N, relative cell viability of CD8 T cells, NK cells, or H22 cells processed with ultrasound irradiation and mixed cultivation in transwells. O-P, relative cell viability of CD8 T cells, NK cells, or Hepa1-6 cells processed with ultrasound irradiation and mixed cultivation in transwells. Q, FCM results of Fluo-4 fluorescent CD8 T cells. C1, H22/Hepa1-6 cancer cells; C2, CD8 T/NK cells; C3, CD8 T/NK cells cocultured with H22/Hepa1-6 cancer cells in the transwell; C4, FUS stimulated CD8 T/NK cells; C5, CD8 T/NK cells subjected to FUS irradiation, and cocultured with H22/Hepa1-6 cancer cells in the transwell; C6, CD8 T/NK cells mixed with H22/Hepa1-6 lysate and cocultured with H22/Hepa1-6 cancer cells in the transwell; C7, CD8 T/NK cells in 300 nM calcium culture medium and cocultured with H22/Hepa1-6 cancer cells in the transwell; C8, CD8 T/NK cells mixed with H22/Hepa1-6 lysate, subjected to FUS irradiation, and cocultured with H22/Hepa1-6 cancer cells in the transwell; C9, CD8 T/NK cells mixed with H22/Hepa1-6 lysate in 300 nM calcium culture medium and cocultured with H22/Hepa1-6 cancer cells in the transwell; C10, CD8 T/NK cells in 300 nM calcium culture medium subjected to FUS irradiation, and cocultured with H22/Hepa1-6 cancer cells in the transwell; C11, CD8 T/NK cells mixed with H22/Hepa1-6 lysate in 300 nM calcium culture medium, subjected to FUS irradiation, and cocultured with H22/Hepa1-6 cancer cells in the transwell.

Interestingly, calcium content assay, and FCM of Fluo-4 AM labeled cells consistently proved a significant increase of calcium in the splenocytes after FUS sti. spleen (Figure 5 I-J). Furthermore, qRT-PCR results showed a significant upregulation of classic genes related to TNF, NFκB, MAPK, and other signaling pathways (Figure 5 K-L). Note that although FUS sti. spleen significantly upregulated PIEZO1 expression, genes associated with calcium channels, such as TRPV, TRPC, TRPA, or TRPM, did not change in response to ultrasound (Figure 5 K-L). Building on previous studies showing that ultrasound facilitating calcium influx [98, 99], we speculated that the ultrasonic mechanical force, microjet, vaporization, or cavitation effects generated in the splenic microenvironment leading to sonoporation in the cell membrane, thus directly promoting calcium inflow through the membrane gaps but not eliciting extensive changes in cellular ion channels (such as TRP channels) [98, 99]. This aspect lies beyond the current research scope and merits independent investigation in future studies.

To verify the ability of FUS to increase Ca^2+^ influx that modulated anticancer immunocompetence, CD8 T cells (CD45^+^ CD3^+^ CD8a^+^) and NK cells (CD45^+^ CD3^-^ NK1.1^+^) were sorted, subjected to ultrasound irradiation, and then cocultured with cancer cells in transwell. The viability of both CD8 T cells and NK cells was significantly higher than that of other groups after ultrasound irradiation especially under the condition of high calcium, and more importantly, the H22 cancer cell viability was significantly lower (Figure 5 M-N). Additionally, the results of repeated experiments with GFP-labeled Hepa1-6 cancer cells further proved that ultrasound stimulation significantly activated CD8 T cells or NK cells immunocompetency to suppress Hepa1-6 cancer cell proliferation especially under high calcium (Figure 5 O-P; Supplementary Figure 13). Moreover, FCM results of Fluo-4 fluorescent cells further demonstrated that ultrasound-mediated calcium influx significantly enhances immune cell cytotoxicity, thereby potentiating the tumoricidal activity (Figure 5 Q). However, the FUS exposure remarkably highlighted immune cell immunocompetence to suppress the proliferation of cancer cells only under the co-stimulation of cancer cell lysate (denoting tumor cytokines or cancer cell antigens) (Figure 5 M-P).

### 3.4 FUS plus STNDs@Ca^2+^ activated splenic immunity to suppress orthotopic H22-HCC proliferation

Intracellular Ca^2+^ are intricately involved in a multitude of signaling pathways, yet there remains a significant gap in the development of effective methodologies to selectively inhibit calcium signaling within specific pathways. Under ultrasonic conditions, the pathways through which Ca^2+^ enters cells are notably diverse. While the preliminary findings in section 3.3 have demonstrated that ultrasound-induced cell membrane perforation facilitates Ca^2+^ influx, there is a conspicuous absence of targeted strategies to inhibit the subsequent biophysical processes following this influx.

This limitation stems from the inherent complexity of calcium dynamics, where the spatial and temporal resolution of current techniques is insufficient to precisely delineate the source and destination of calcium signals within the cellular milieu. Consequently, this study refrains from exploring the intricate regulatory mechanisms of specific genes or signaling pathways mediated by Ca^2+^.

The primary objective of this research is to explore the methodologies and therapeutic efficacy of ultrasound in modulating spleen-mediated immune suppression of cancer. Building upon the aforementioned findings that ultrasound significantly enhances Ca^2+^ influx, thereby potentiating the spleen’s intrinsic capacity to suppress tumorigenesis, and taking into account the physiologically constrained calcium levels within the splenic microenvironment, we hypothesized that targeted pharmacological augmentation of calcium availability in the spleen could synergistically amplify its anti-neoplastic efficacy. To address this hypothesis, we innovatively designed the spleen-targeted nanodroplets loading Ca^2+^ (STNDs@Ca^2+^; Figure 6 A), which are engineered to undergo phase transition and cavitation upon FUS exposure, thereby facilitating the targeted and controlled release of Ca^2+^ to splenic immunocytes (Figure 6 B). During this process, the sonoporation induced by FUS plus STNDs@Ca^2+^ significantly surpasses that of FUS alone in both intensity and frequency of disrupting the cellular membrane barrier. This enhanced physical disruption provides a greater number of opportunities for calcium influx, thereby facilitating a more pronounced intracellular calcium elevation (Figure 6 B).

**Figure 6.**
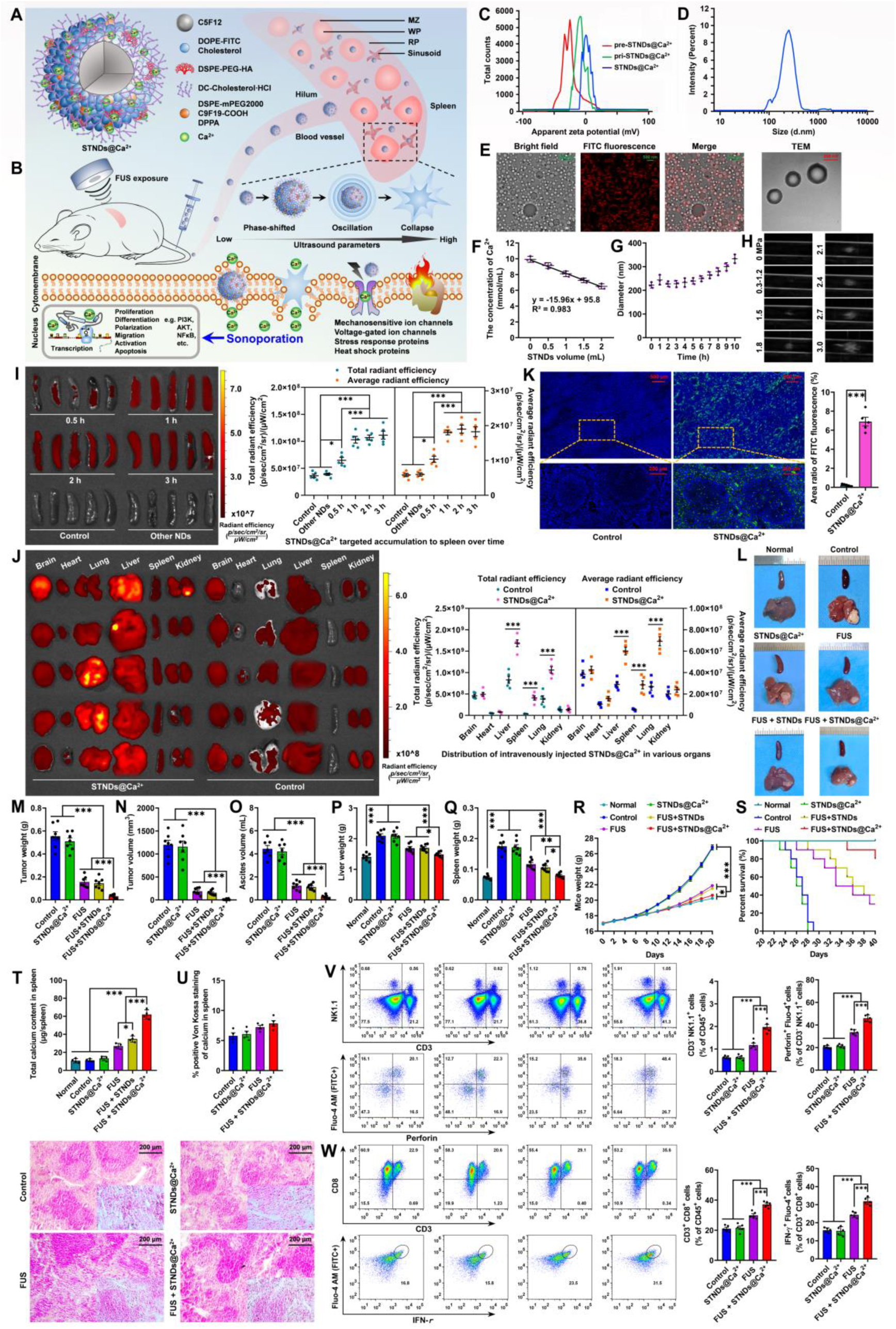
Characterization of STNDs@Ca^2+^ and therapeutic efficacy evaluation of FUS plus STNDs@Ca^2+^. A, structural diagram of the STNDs@Ca^2+^. B, the process and mechanism of FUS plus STNDs@Ca^2+^ facilitated Ca^2+^ controlled-release in the spleen, leading to enhanced immune cell activation and antitumor effects. C, zeta potential changes during the preparation of STNDs@Ca^2+^, showing a transition from negative to neutral. D, size distribution of STNDs@Ca^2+^. E, LCFM and cryo-TEM images demonstrating spherical morphology and uniform size (∼247 ± 23 nm). F, calcium-loading capacity of STNDs@Ca^2+^, measured at ∼2 mmol per 1 mL STNDs@Ca^2+^. G, dynamic changes in STNDs@Ca^2+^ size over time at 37°C. H, B-mode ultrasound imaging of FUS-triggered phase transition and cavitation of STNDs@Ca^2+^ under varying peak negative pressures. I, ILLIS monitoring splenic accumulation of the STNDs@Ca^2+^ within the 0-3 hour time window. J, ILLIS imaging of in vivo biodistribution of STNDs@Ca^2+^ after intravenous injection. K, microscope images showing fluorescent STNDs@Ca^2+^ distribution in spleen and the corresponding positive area ratio analysis. L-S, therapeutic outcomes in orthotopic H22-HCC mice, including anatomical pictures, tumor weight, tumor volume, ascites volume, liver weight, spleen weight, mice weight and survival curves. T, total calcium content in the spleen. U, microscope images of Von Koussa staining and the corresponding positive area ratio analysis. V-W, FCM analysis of the proportions of NK cells and CD8^+^ T cells, particularly calcium-positive activated subsets.

During the fabrication of STNDs@Ca^2+^, the zeta potential (Figure 6 C) of the precursors, intermediate products, and final formulation transitioned from a negative value (-26.4 ± 3.2 mV) to a near-neutral value (-0.7 ± 2.6 mV), indicating successful surface modification and calcium incorporation. Dynamic light scattering (DLS) analysis revealed that the STNDs@Ca^2+^ exhibited a uniform hydrodynamic diameter of approximately 250 nm (Figure 6 D), with a polydispersity index (PDI) of ∼0.3, indicating a narrow size distribution. High-resolution imaging techniques, including LCFM and cryo-TEM, further confirmed the spherical morphology and monodispersity of the nanodroplets (Figure 6 E), with an average diameter consistent with DLS measurements. The calcium encapsulation efficiency of the nanodroplets was quantified through calcium colorimetric assay, revealing a loading capacity of approximately 2 mmol Ca²⁺ per 1 mL nanodroplets (Figure 6 F). Furthermore, the STNDs@Ca^2+^ maintained excellent stability for up to 5 h at 37℃, with their size consistently remaining around 250 nm (Figure 6 G). B-mode ultrasound imaging revealed that the nanodroplets underwent phase transition and cavitation at a threshold acoustic pressure of 1.5 MPa (Figure 6 H). As the peak negative pressure increased, the cavitation effects became more pronounced with enhanced echogenicity and bubble formation. This phenomenon confirms the potential of STNDs@Ca^2+^ for controlled release of calcium ions. Based on the preliminary experimental findings, we strategically selected an acoustic pressure of 2.1 MPa for subsequent investigations to produce the desired physical effects without potential tissue damage.

ILLIS results demonstrated that following intravenous administration of 200 μL STNDs@Ca^2+^ in mice, continuous monitoring within the 0-3 hour time window revealed a plateau in splenic accumulation of the STNDs@Ca^2+^ at 1 hour post-administration, with no significant variation in fluorescence intensity observed thereafter. This pharmacokinetic profile suggests that the nanodroplets reached a distribution equilibrium in the spleen within 1 hour of administration (Figure 6 I). Additionally, quantitative biodistribution analysis revealed significant accumulation in the liver, lungs, and spleen, demonstrating the nanoparticles’ preferential tropism towards these reticuloendothelial system-rich organs (Figure 6 J). Moreover, microscopic imaging further confirmed the effective accumulation of STNDs@Ca^2+^ targeting the spleen (Figure 6 K). This tissue distribution pattern is consistent with the physiological characteristics of nanoparticle clearance and the vascular architecture of these organs. Importantly, comprehensive toxicological evaluation, including serum biochemical profiling (Table 2) and histopathological examination (Supplementary Figure 14 A) of hepatic, pulmonary, and splenic tissues, demonstrated an absence of toxicity. Histological sections showed preserved tissue architecture without evidence of inflammatory infiltration, cellular necrosis, or other pathological alterations, while serum markers remained within normal physiological ranges. These findings collectively indicate that the STNDs@Ca^2+^ exhibit favorable biocompatibility and safety profile at the administered dose, supporting their potential for therapeutic applications.

**Table 2.** The results of serum biochemical analysis to evaluate the toxicity of STNDs@Ca^2+^. O1, the saline injected group; O2, the STNDs@Ca^2+^ injected group.

| Biochemical analysis | O1 (Average ± SEM) | O2 (Average ± SEM) |
| --- | --- | --- |
| ALT (U/L) | 94.53 ± 4.82 | 90.20 ± 3.17 |
| AST (U/L) | 247.51 ± 8.35 | 253.62 ± 10.72 |
| TBIL (μmol/L) | 13.52 ± 1.46 | 14.68 ± 2.09 |
| UREA (mg/dl) | 26.83 ± 2.34 | 24.37 ± 1.63 |
| CREA (μmol/L) | 28.14 ± 2.31 | 32.28 ± 3.36 |
| UA (μmol/L) | 124.27 ± 9.54 | 132.49 ± 11.38 |
| CK (U/L) | 2422.64 ± 112.58 | 2538.46 ± 96.14 |

The deteriorated H22-HCC was commonly accompanied by malignant ascites that were a common and distressing condition associated with an advanced stage of neoplasms, typically connected to a poor prognosis and a median survival time of no more than 4 months in clinical practice, which was more consistent with the characteristics of HCC in humans and probably the reasonable model for clinically evaluating HCC therapy [100–103]. Therefore, to evaluate the therapeutic potential of FUS plus STNDs@Ca^2+^, experimental mice were randomly divided into six experimental groups: S1 group with normal mice intravenously injected with saline classified as negative control (named normal), S2 group with orthotopic H22-HCC mice administered with saline used as the positive control (named control), S3 group with orthotopic H22-HCC mice intravenously injected with STNDs@Ca^2+^ (named STNDs@Ca^2+^), S4 group with orthotopic H22-HCC mice administered with saline and subjected to SUS (named FUS sti. spleen), S5 group with orthotopic H22-HCC mice subjected to FUS plus STNDs (named FUS + STNDs), and S6 group with orthotopic H22-HCC mice subjected to FUS plus STNDs@Ca^2+^ (named FUS + STNDs@Ca^2+^). The FUS + STNDs@Ca^2+^ group demonstrated the most pronounced antitumor effects (Figure 6 L-N; Supplementary Figure 15), with a tumor inhibition rate of approximately 93%, significantly outperforming the FUS sti. spleen and FUS + STNDs groups (with a tumor suppression rate of about 69%). Additionally, the FUS + STNDs@Ca^2+^ group exhibited a remarkable ascites inhibition rate of over 90%, compared to 70% in the other FUS-treated groups (Figure 6 O). Organ and body weight analysis further corroborated these findings, with the FUS + STNDs@Ca^2+^ group showing significantly reduced liver and spleen weights than other groups, indicative of attenuated tumor burden and splenomegaly (Figure 6 P-R). Survival analysis revealed a substantial extension in the lifespan of orthotopic H22-HCC mice treated with FUS plus STNDs@Ca^2+^ (Figure 6 S), highlighting the therapeutic superiority of this approach. Quantitative analysis of total calcium content in the spleen revealed significantly higher levels in the FUS + STNDs@Ca^2+^ group compared to the others (Figure 6 T). Ultrasound-mediated calcium influx may lead to minimal calcium deposition; however, no statistically significant difference was observed compared to the control group (Figure 6 U). Notably, accumulating evidence has demonstrated that physiological levels of calcium deposition (calcification) can potentiate immune system activation, particularly by enhancing the immunoregulatory functions of macrophages [104–108]. Additionally, histological assessment through HE and TUNEL assays revealed no significant pathological alterations or abnormal apoptotic activity in the experimental groups (Supplementary Figure 14 B). Comparison of S4-S6 groups showed that ultrasonic cavitation facilitated greater calcium influx than ultrasound alone, especially with exogenous calcium supplementation, which significantly enhanced intracellular calcium levels (Figure 6 T). Moreover, FCM analysis demonstrated a marked increase in the proportion of NK cells and CD8⁺ T cells in the FUS + STNDs@Ca^2+^ group, of which the proportion of activated (Ca²⁺-positive) NK and CD8⁺ T cells was significantly higher than that of the other groups (Figure 6 V-W), further supporting the role of Ca^2+^ in immune cell activation and antitumor immunity. Notably, the ultrasound-mediated calcium influx was not specifically targeted to any particular cellular subtype or population, as demonstrated by heterogeneous immunomodulatory patterns and conserved enrichment profiles of key signaling pathways in differentially expressed genes among splenic T/NK cells, myeloid lineages, and B cell subsets (Section 3.2; Supplementary File 1), suggesting calcium-dependent immunoregulation operates through conserved mechanistic pathways across multiple immune cell lineages.

## 4 Discussion

### 4.1 FUS sti. spleen reduced splenic and intratumoral MDSCs

MDSCs are a heterogeneous population of neutrophil and monocyte-like myeloid cells, which are generally considered as important mediators of immune suppression, angiogenesis, tumor invasion, and metastatic progression in various types of tumor [109]. The unique transcriptional signatures of MDSCs are currently confused, and no recognized signature markers are available to clearly delineate unique subpopulations of MDSCs to distinguish them from neutrophils and monocytes [109]. Constrainedly, we applied the emblematic markers Itgam/Ly6c2/Ly6g to classify the splenic/intratumoral PMN-MDSCs and M-MDSCs, respectively, as reported by Alshetaiwi H er al. [109] and Bronte V et al. [110]. Besides, Cd84 usually participates in cell-cell interactions and modulation of the activation and differentiation of a variety of immune cells, particularly regulating the intratumoral immunosuppressive microenvironment as a suppressor of T cell activation [111, 112]. Recent research by Alshetaiwi H et al. [109] demonstrated that Cd84 was significantly upregulated in MDSCs, which exerted a significant T cell suppression, thus identifying Cd84 as a robust and generalizable cell surface marker for PMN- and M-MDSCs. Accordingly, in this study, PMN-MDSCs were defined as Itgam^+^ Ly6c2^low^ Ly6g^+^ Cd84^high^ and M-MDSCs as Itgam^+^ Ly6c2^+^ Ly6g^-^ Cd84^high^. However, these markers clustering distinct MDSC subgroups still overlapped with those defined myeloid counterparts (Supplementary Figure 16 A-B), suggesting that a similar immune-suppressive cell state can be acquired by both monocytes and neutrophils independently, as stated in the Supplementary File 1. FUS sti. spleen elicited a dual immunomodulatory effect, characterized by a substantial reduction in both PMN-MDSC and M-MDSC populations across both the splenic and tumoral microenvironments (Supplementary Figure 16 C), suggesting a significant attenuation of immunosuppressive microenvironment within both the spleen and tumor niche (Supplementary Figure 16 D). Itgam^+^ CD14^+^ and Itgam^+^ Fut4^+^ were also used to define the PMN- and M-MDSCs, respectively, as previous reported [110, 113]. In general, Itgam/CD14/Fut4 markers were also expressed in both monocytes and neutrophils, and in particular, the cell proportion of PMN- and M-MDSCs in the spleen and tumor from FUS sti. spleen group was significantly lower than that of the control group (Supplementary Figure 16 E-H), consistent with the above results. The combined results of scRNA-seq and FCM consistently demonstrated that FUS sti. spleen significantly reduced the proportion of MDSCs, suggesting that negative immunity was suppressed. Besides, the concomitant downregulation of GM-CSF coupled with the upregulation of IL-1β provides corroborative evidence for the diminished immunosuppressive capacity of MDSCs (Figure 4 S) [109].

### 4.2 Ultrasound facilitated Ca^2+^ influx potentiating immunocyte immunocompetence

Calcium is indispensable secondary messenger that orchestrate a myriad of cellular processes, including proliferation, activation, polarization, migration, and metabolism, across various cell types [114, 115]. Particularly, in the context of immunology, Ca^2+^ signaling is integral to the immunocyte immunocompetence against assorted illnesses. For instance, in T cells, Ca^2+^ influx through CRAC (calcium release-activated calcium), TRP (transient receptor potential), and voltage-gated channels is essential for T cell receptor (TCR) signaling, leading to the activation of transcription factors such as NFAT (nuclear factor of activated T cells), which drive cell proliferation, differentiation, cytokine secretion and cytolytic cell function [116–120]. Similarly, in NK cells, Ca^2+^ signaling is critical for the formation of the immunological synapse and the subsequent release of cytotoxic granules containing perforin and granzymes, which are crucial for target cell killing, underscoring the therapeutic potential of targeting Ca^2+^ signaling pathways [121–124]. Additionally, the other immunocyte phenotypes, including CD4 T cell, B cell, monocyte/Mφ, DC, and neutrophil subsets, are also governed by Ca^2+^ signaling, which significantly functioned in innate and adaptive immunity to maintain host immune homeostasis [115]. Thus, Ca^2+^ serves as a master regulator of immune cell function, making it a compelling target for therapeutic intervention, particularly for tumor immunotherapy [115, 118, 125].

This study convincingly evidenced that FUS sti. spleen potentiates calcium influx to modulate multiple immunoregulatory signaling cascades, including but not limited to TNF, NFκB, MAPK, HIF-1, and ErbB pathways, across heterogeneous immunocyte populations, particularly CD8+ T lymphocytes and NK cells, thereby counteracting tumor-driven immunosuppressive polarization while potentiating robust splenic immune activation that impedes malignant progression and neoplastic proliferation. Leveraging these mechanistic insights, we developed an innovative acoustically responsive STNDs@Ca^2+^ nanocomposite that undergoes ultrasound-triggered phase-transition and cavitation to control exogenous Ca^2+^ release within the splenic microenvironment, thereby potentiating more superior splenic immunomodulatory efficacy and augmenting better antitumor therapeutic outcomes compared to ultrasound monotherapy (Figure 6). Despite the current investigation encountering methodological constraints in delineating the hierarchical calcium signaling network and further identifying the apex molecular regulator, which represents a transformative therapeutic frontier whose strategic manipulation, either as a monotherapeutic strategy or in synergistic combination with exogenous calcium delivery, may establish a transformative paradigm for immunopotentiation-based cancer therapy, warranting comprehensive mechanistic investigation in future research endeavors.

Indeed, substantial evidence demonstrated that diverse methodologies, particularly pH-responsive and other functionalized calcium-loaded nanoparticles, modulating calcium influx in immune cells have been successfully implemented as promising therapeutic strategies for various pathologies, including neoplastic diseases [126–129]. Among that, ultrasound has revolutionized this paradigm by offering a non-invasive nature, precise spatial targeting capability, exceptional tissue penetrability, and controllable modality for targeted calcium ion modulation [96, 130–134]. While only a handful of studies preliminary evidenced that low-intensity ultrasound facilitated calcium influx through mechanosensitive ion channels, notably Piezo1, triggering activation of specialized immunocyte subsets [135, 136], no previous investigation has systematically demonstrated the therapeutic potential of ultrasound-mediated calcium modulation in orchestrating splenic immune responses or non-specific immunomodulatory effects across diverse immunocyte populations for disease treatment, particularly in oncological applications. It is crucial to emphasize that under identical acoustic parameters, the mechanical forces and cell perforation generated by ultrasound alone or in combination with endogenous cavitation nuclei are significantly inferior in magnitude, intensity, and frequency compared to those elicited by ultrasound coupled with artificial cavitation nuclei, such as microbubbles, nanobubbles, and phase-shifted nanodroplets [131, 137–141]. Nevertheless, meticulous optimization of acoustic parameters is imperative to mitigate potential tissue damage while ensuring optimal therapeutic efficacy, as evidenced by differential cellular responses under varying acoustic pressures where FUS at 2.4 MPa alone caused no cellular damage but triggered apoptosis in the presence of STNDs@Ca²⁺, whereas no significant cell damage was observed at 2.1 MPa (Supplementary Figure 14 B). Moreover, artificial cavitation nuclei exhibit remarkable design flexibility, as exemplified by the STNDs@Ca^2+^ developed in this study, which demonstrates dual functionality of calcium encapsulation and splenic targeting, thereby facilitating localized mechanotransduction and potentiates calcium influx in nearby immunocyte. The STNDs@Ca^2+^ fabrication was accomplished through a strategic modification on superficial phospholipid composition of conventional nanodroplets, leveraging the spleen-targeting affinity of DOPE and cholesterol [142–144], to enable spleen-specific accumulation, while Ca^2+^ encapsulation was achieved via an electro-potential coupling mechanism [145, 146]. Future investigations may explore the strategic functionalization of STNDs@Ca^2+^ conjugated with ligands, guided by the distinct splenic architecture and splenocyte compartmentalization, to optimize therapeutic precision and efficacy through spatially controlled calcium delivery to specific immunocyte populations. In summary, the integration of ultrasound with immunotherapy and nanotechnology holds immense potential for revolutionizing cancer treatment and other immune-related disorders. However, further studies are needed to fully understand the mechanisms involved and to translate these findings into clinical practice.

## 5 Conclusion

In conclusion, this study elucidated the profound therapeutic potential of ultrasound in modulating splenic immunity to combat HCC. The comprehensive analysis of the dynamic immune landscape revealed that FUS sti. spleen not only attenuated tumor-induced immunosuppression but also revitalized the antitumor capabilities of splenic immunocytes, particularly CD8^+^ T cells and NK cells. The mechanistic insights revealed that FUS-facilitated calcium influx played a pivotal role in activating intracellular signaling pathways such as TNF, NFκB, MAPK, and HIF-1, which collectively promoted splenic immunocyte activation and migration to the tumor microenvironment, thereby significantly suppressing tumor proliferation and improving survival outcomes in murine models of H22-HCC and Hepa1-6-HCC. These findings underscored the critical role of the spleen in regulating systemic antitumor immunity and demonstrate the ability of FUS to reprogram the immune landscape in favor of tumor suppression.

Furthermore, FUS triggering the innovative STNDs@Ca²⁺ cavitation enabled controlled-release of exogenous calcium to splenic immunocytes, amplifying the immunostimulatory effects and achieving superior tumor suppression (such as a remarkable H22-HCC suppression rate of up to 93%) compared to FUS alone. Such synergistic approach not only enhanced the specificity and efficacy of the treatment but also minimizes potential off-target effects, underscoring its translational potential for clinical applications.

By leveraging the biophysical properties of ultrasound and the immunogenic potential of calcium signaling, this work opens new avenues for the development of noninvasive, targeted immunotherapies. Future studies should focus on optimizing the delivery and specificity of calcium-loaded nanodroplets, as well as exploring the broader applicability of this approach in other cancer types and immune-related disorders. Ultimately, this work paves the way for a paradigm shift in cancer immunotherapy, offering foundation for the development of noninvasive, precision-based immunotherapies.

## 6 Abbreviations

FUS, focused ultrasouond; H22, Hepatoma-22; HCC, hepatocellular carcinoma; SUS, splenic ultrasound stimulation; TME, tumor microenvironment; FUS sti. spleen, FUS noninvasively stimulated the spleen; FCM, flow cytometry; UMAP, Uniform Manifold Approximation and Projection; DEGs, differentially expressed genes; CV, crystal violet staining; GSVA, gene set variation analysis; GSEA, gene set enrichment analysis; TCR, T cell receptor; scRNA-seq, single-cell RNA sequencing; CNV, copy number variation; bRNA-seq, bulk RNA sequencing; DCs, dendritic cells; Treg, regulatory T cells; NK cells, natural killer cells; M- and PMN-MDSCs, (monocytic and polymorphonuclear) myeloid-derived suppressor cells; Mφ, macrophage; MSCs, myeloid stem cells; KEGG, Kyoto Encyclopedia of Genes and Genomes; GO, Gene Ontology; STNDs@Ca^2+^, spleen-targeted nanodroplets encapsulating bioavailable calcium ions.

## 7 Ethical approval

The experiment was approved by the Animal Ethics Committee of Xi’an Jiaotong University before the research. The number of the ethics approval was XJTU_2022-0033. The animal experiments were conducted following ARRIVE guidelines. All animals had access to water and food *ad libitum* and were maintained in a pathogen-free facility at 25 ± 1 °C and 40 ± 10% relative humidity under a 12-h light/dark cycle. At the end of study, the animals were sacrificed with cervical dislocation method following anesthesia with 1.5% isoflurane.

## 8 Financial supports

This work was supported by the National Natural Science Foundation of China (No. 12204370), the Innovation Ability Supporting Program of Shaanxi Province (No. 2023WGZJ-ZD-09), the Basic-Clinical Integration Innovation Project of Xi’an Jiaotong University (No. YXJLRH2022092), and the Natural Science Basic Research Program of Shaanxi Province (No. 2023-JC-QN-0036).

## 10 Authors’ contributions

Wei Dong performed the animal experiments, FCM analysis, scRNA-seq & bRNA-seq analysis, IHC staining assays, and wrote the manuscript. Guihu Wang and Senyang Li performed the IHC staining assays and in vitro experiments. Other authors also performed in vivo and in vitro experiments. Pengfei Liu and Zongfang Li interpreted the data and revised the manuscript. All authors designed the experiments, interpreted the data, provided supervision, wrote the manuscript, and approved the submitted version.

## Acknowledgements

We thank Prof. Guangyao Kong, Prof. Shemin Lv, and Prof. Tielin Yang for their valuable suggestions. We are very grateful for the technical support of scRNA-seq, scTCR-seq, bRNA-seq, and bioinformatics analysis from OE Biotech Co., Ltd. (Shanghai, China). We would like to thank MogoEdit (https://www.mogoedit.com) for its English editing during the preparation of this manuscript.

## 11 Supplementary data

The supplementary material for this article can be found in the Supplementary Figures and Tables.

## 12 Data availability statement

The scRNA-seq, TCR-seq, and bRNA-seq data presented in the study are deposited in the NCBI Gene Expression Omnibus database (https://www.ncbi.nlm.nih.gov/), accession number GSE277240, GSE267237, and GSE267445.

## Reference

[1] W.H. Crosby, An historical sketch of splenic function and splenectomy, Lymphology 16(2) (1983) 52–5.

[2] R.J. Holdsworth, A.D. Irving, A. Cuschieri, Postsplenectomy sepsis and its mortality rate: actual versus perceived risks, Br J Surg 78(9) (1991) 1031–8.

[3] N. Bisharat, H. Omari, I. Lavi, R. Raz, Risk of infection and death among post-splenectomy patients, J Infect 43(3) (2001) 182–6.

[4] J. Higashijima, M. Shimada, M. Chikakiyo, T. Miyatani, K. Yoshikawa, M. Nishioka, T. Iwata, N. Kurita, Effect of splenectomy on antitumor immune system in mice, Anticancer Res 29(1) (2009) 385–93.

[5] T. Toge, K. Kuroi, H. Kuninobu, Y. Yamaguchi, Y. Kegoya, N. Baba, T. Hattori, Role of the spleen in immunosuppression of gastric cancer: predominance of suppressor precursor and suppressor inducer T cells in the recirculating spleen cells, Clin Exp Immunol 74(3) (1988) 409–12.

[6] B. Li, S. Zhang, N. Huang, H. Chen, P. Wang, J. Li, Y. Pu, J. Yang, Z. Li, Dynamics of the spleen and its significance in a murine H22 orthotopic hepatoma model, Exp Biol Med (Maywood) 241(8) (2016) 863–72.

[7] F.A. Koopman, S.S. Chavan, S. Miljko, S. Grazio, S. Sokolovic, P.R. Schuurman, A.D. Mehta, Y.A. Levine, M. Faltys, R. Zitnik, K.J. Tracey, P.P. Tak, Vagus nerve stimulation inhibits cytokine production and attenuates disease severity in rheumatoid arthritis, Proc Natl Acad Sci U S A 113(29) (2016) 8284–9.

[8] E.M. Annoni, X. Xie, S.W. Lee, I. Libbus, B.H. KenKnight, J.W. Osborn, E.G. Tolkacheva, Intermittent electrical stimulation of the right cervical vagus nerve in salt-sensitive hypertensive rats: effects on blood pressure, arrhythmias, and ventricular electrophysiology, Physiol Rep 3(8) (2015).

[9] D.W. Wang, Y.M. Yin, Y.M. Yao, Vagal Modulation of the Inflammatory Response in Sepsis, Int Rev Immunol 35(5) (2016) 415–433.

[10] T. Inoue, C. Abe, S.S. Sung, S. Moscalu, J. Jankowski, L. Huang, H. Ye, D.L. Rosin, P.G. Guyenet, M.D. Okusa, Vagus nerve stimulation mediates protection from kidney ischemia-reperfusion injury through α7nAChR+ splenocytes, J Clin Invest 126(5) (2016) 1939–52.

[11] B. Bonaz, V. Sinniger, D. Hoffmann, D. Clarençon, N. Mathieu, C. Dantzer, L. Vercueil, C. Picq, C. Trocmé, P. Faure, J.L. Cracowski, S. Pellissier, Chronic vagus nerve stimulation in Crohn’s disease: a 6-month follow-up pilot study, Neurogastroenterol Motil 28(6) (2016) 948–53.

[12] B. Bonaz, V. Sinniger, S. Pellissier, Vagus nerve stimulation: a new promising therapeutic tool in inflammatory bowel disease, J Intern Med 282(1) (2017) 46–63.

[13] S.F. Cogan, Neural stimulation and recording electrodes, Annual review of biomedical engineering 10 (2008) 275–309.

[14] D.P. Zachs, S.J. Offutt, R.S. Graham, Y. Kim, J. Mueller, J.L. Auger, N.J. Schuldt, C.R.W. Kaiser, A.P. Heiller, R. Dutta, H. Guo, J.K. Alford, B.A. Binstadt, H.H. Lim, Noninvasive ultrasound stimulation of the spleen to treat inflammatory arthritis, Nat Commun 10(1) (2019) 951.

[15] S. Wang, W. Meng, Z. Ren, B. Li, T. Zhu, H. Chen, Z. Wang, B. He, D. Zhao, H. Jiang, Ultrasonic Neuromodulation and Sonogenetics: A New Era for Neural Modulation, Frontiers in physiology 11 (2020) 787.

[16] C.T. Curley, N.D. Sheybani, T.N. Bullock, R.J. Price, Focused Ultrasound Immunotherapy for Central Nervous System Pathologies: Challenges and Opportunities, Theranostics 7(15) (2017) 3608–3623.

[17] B. Chertok, R. Langer, Circulating Magnetic Microbubbles for Localized Real-Time Control of Drug Delivery by Ultrasonography-Guided Magnetic Targeting and Ultrasound, Theranostics 8(2) (2018) 341–357.

[18] A. Lakshmanan, Z. Jin, Acoustic biosensors for ultrasound imaging of enzyme activity, 16(9) (2020) 988–996.

[19] M.D. Okusa, D.L. Rosin, K.J. Tracey, Targeting neural reflex circuits in immunity to treat kidney disease, Nature reviews. Nephrology 13(11) (2017) 669–680.

[20] V. Cotero, Y. Fan, T. Tsaava, A.M. Kressel, I. Hancu, P. Fitzgerald, K. Wallace, S. Kaanumalle, J. Graf, W. Rigby, T.J. Kao, J. Roberts, C. Bhushan, S. Joel, T.R. Coleman, S. Zanos, K.J. Tracey, J. Ashe, S.S. Chavan, C. Puleo, Noninvasive sub-organ ultrasound stimulation for targeted neuromodulation, Nat Commun 10(1) (2019) 952.

[21] J.C. Gigliotti, L. Huang, H. Ye, A. Bajwa, K. Chattrabhuti, S. Lee, A.L. Klibanov, K. Kalantari, D.L. Rosin, M.D. Okusa, Ultrasound prevents renal ischemia-reperfusion injury by stimulating the splenic cholinergic anti-inflammatory pathway, J. Am. Soc. Nephrol. 24(9) (2013) 1451–60.

[22] U. Ahmed, J.F. Graf, A. Daytz, O. Yaipen, I. Mughrabi, N. Jayaprakash, V. Cotero, C. Morton, C.S. Deutschman, S. Zanos, C. Puleo, Ultrasound Neuromodulation of the Spleen Has Time-Dependent Anti-Inflammatory Effect in a Pneumonia Model, Front Immunol 13 (2022) 892086.

[23] J.C. Gigliotti, L. Huang, H. Ye, A. Bajwa, K. Chattrabhuti, S. Lee, A.L. Klibanov, K. Kalantari, D.L. Rosin, M.D. Okusa, Ultrasound prevents renal ischemia-reperfusion injury by stimulating the splenic cholinergic anti-inflammatory pathway, J Am Soc Nephrol 24(9) (2013) 1451–60.

[24] X. Hu, F. Li, J. Zeng, Z. Zhou, Z. Wang, J. Chen, D. Cao, Y. Hong, L. Huang, Y. Chen, J. Xu, F. Dong, R. Yu, H. Zheng, Noninvasive Low-Frequency Pulsed Focused Ultrasound Therapy for Rheumatoid Arthritis in Mice, Research 2022 (2022) 0013.

[25] E. Memari, D. Khan, R. Alkins, B. Helfield, Focused ultrasound-assisted delivery of immunomodulating agents in brain cancer, J. Control Release 367 (2024) 283–299.

[26] F. Ivanovski, M. Meško, T. Lebar, M. Rupnik, D. Lainšček, M. Gradišek, R. Jerala, M. Benčina, Ultrasound-mediated spatial and temporal control of engineered cells in vivo, Nat. Commun. 15(1) (2024) 7369.

[27] D.P. Zachs, S.J. Offutt, R.S. Graham, Y. Kim, J. Mueller, J.L. Auger, N.J. Schuldt, C.R.W. Kaiser, A.P. Heiller, R. Dutta, H. Guo, J.K. Alford, B.A. Binstadt, H.H. Lim, Noninvasive ultrasound stimulation of the spleen to treat inflammatory arthritis, Nat. Commun. 10(1) (2019) 951.

[28] A. Butler, P. Hoffman, P. Smibert, E. Papalexi, R. Satija, Integrating single-cell transcriptomic data across different conditions, technologies, and species, Nat Biotechnol 36(5) (2018) 411–420.

[29] L. Haghverdi, A.T.L. Lun, M.D. Morgan, J.C. Marioni, Batch effects in single-cell RNA-sequencing data are corrected by matching mutual nearest neighbors, Nat Biotechnol 36(5) (2018) 421–427.

[30] X. Qiu, A. Hill, J. Packer, D. Lin, Y.A. Ma, C. Trapnell, Single-cell mRNA quantification and differential analysis with Census, Nat Methods 14(3) (2017) 309–315.

[31] S.V. Puram, I. Tirosh, A.S. Parikh, A.P. Patel, K. Yizhak, S. Gillespie, C. Rodman, C.L. Luo, E.A. Mroz, K.S. Emerick, D.G. Deschler, M.A. Varvares, R. Mylvaganam, O. Rozenblatt-Rosen, J.W. Rocco, W.C. Faquin, D.T. Lin, A. Regev, B.E. Bernstein, Single-Cell Transcriptomic Analysis of Primary and Metastatic Tumor Ecosystems in Head and Neck Cancer, Cell 171(7) (2017) 1611–1624.e24.

[32] V.Y. Kiselev, K. Kirschner, M.T. Schaub, T. Andrews, A. Yiu, T. Chandra, K.N. Natarajan, W. Reik, M. Barahona, A.R. Green, M. Hemberg, SC3: consensus clustering of single-cell RNA-seq data, Nat Methods 14(5) (2017) 483–486.

[33] Y. Hao, B. Miraghazadeh, R. Chand, A.R. Davies, C. Cardinez, K. Kwong, M.B. Downes, R.A. Sweet, P.F. Cañete, L.J. D’Orsogna, D.A. Fulcher, S. Choo, D. Yip, G. Peters, S. Yip, M.J. Witney, M. Nekrasov, Z.P. Feng, D.C. Tscharke, C.G. Vinuesa, M.C. Cook, CTLA4 protects against maladaptive cytotoxicity during the differentiation of effector and follicular CD4(+) T cells, Cell Mol Immunol 20(7) (2023) 777–793.

[34] Y. Kidani, Y. Kitagawa, M. Hagiwara, A. Kawashima, T. Kanazawa, H. Wada, M. Uemura, N. Nonomura, D. Motooka, S. Nakamura, N. Ohkura, S. Sakaguchi, Downregulation of TCF7 and LEF1 is a key determinant of tumor-infiltrating regulatory T-cell function, Int Immunol 36(4) (2024) 167–182.

[35] P.T. Le, N. Ha, N.K. Tran, A.G. Newman, K.M. Esselen, J.L. Dalrymple, E.M. Schmelz, A. Bhandoola, H.H. Xue, P.B. Singh, T.H. Thai, Targeting Cbx3/HP1γ Induces LEF-1 and IL-21R to Promote Tumor-Infiltrating CD8 T-Cell Persistence, Front Immunol 12 (2021) 738958.

[36] B. Petersen, R. Kammerer, A. Frenzel, P. Hassel, T.H. Dau, R. Becker, A. Breithaupt, R.G. Ulrich, A. Lucas-Hahn, G. Meyers, Generation and first characterization of TRDC-knockout pigs lacking γδ T cells, Sci Rep 11(1) (2021) 14965.

[37] Z. Gao, Y. Bai, A. Lin, A. Jiang, C. Zhou, Q. Cheng, Z. Liu, X. Chen, J. Zhang, P. Luo, Gamma delta T-cell-based immune checkpoint therapy: attractive candidate for antitumor treatment, Mol Cancer 22(1) (2023) 31.

[38] R. Förster, A.C. Davalos-Misslitz, A. Rot, CCR7 and its ligands: balancing immunity and tolerance, Nat Rev Immunol 8(5) (2008) 362–71.

[39] D. Dangaj, M. Bruand, A.J. Grimm, C. Ronet, D. Barras, P.A. Duttagupta, E. Lanitis, J. Duraiswamy, J.L. Tanyi, F. Benencia, J. Conejo-Garcia, H.R. Ramay, K.T. Montone, D.J. Powell, Jr., P.A. Gimotty, A. Facciabene, D.G. Jackson, J.S. Weber, S.J. Rodig, S.F. Hodi, L.E. Kandalaft, M. Irving, L. Zhang, P. Foukas, S. Rusakiewicz, M. Delorenzi, G. Coukos, Cooperation between Constitutive and Inducible Chemokines Enables T Cell Engraftment and Immune Attack in Solid Tumors, Cancer Cell 35(6) (2019) 885–900.e10.

[40] J. Zhang, J. Ahn, Y. Suh, S. Hwang, M.E. Davis, K. Lee, Identification of CTLA2A, DEFB29, WFDC15B, SERPINA1F and MUP19 as Novel Tissue-Specific Secretory Factors in Mouse, PLoS One 10(5) (2015) e0124962.

[41] S. Tang, Z. Wu, L. Chen, L. She, W. Zuo, W. Luo, Y. Zhang, S. Liang, G. Liu, B. He, J. He, N. Zhang, Multi-omics analysis reveals the association between elevated KIF18B expression and unfavorable prognosis, immune evasion, and regulatory T cell activation in nasopharyngeal carcinoma, Front Immunol 14 (2023) 1258344.

[42] A. Grandi, M. Tomasi, I. Zanella, L. Ganfini, E. Caproni, L. Fantappiè, C. Irene, L. Frattini, S.J. Isaac, E. König, F. Zerbini, S. Tavarini, C. Sammicheli, F. Giusti, I. Ferlenghi, M. Parri, G. Grandi, Synergistic Protective Activity of Tumor-Specific Epitopes Engineered in Bacterial Outer Membrane Vesicles, Front Oncol 7 (2017) 253.

[43] T.K. Tan, C. Zhang, T. Sanda, Oncogenic transcriptional program driven by TAL1 in T-cell acute lymphoblastic leukemia, Int J Hematol 109(1) (2019) 5–17.

[44] N. Thielemans, S. Demeyer, N. Mentens, O. Gielen, S. Provost, J. Cools, TAL1 cooperates with PI3K/AKT pathway activation in T-cell acute lymphoblastic leukemia, Haematologica 107(10) (2022) 2304–2317.

[45] P.C. Saether, I.H. Westgaard, L.M. Flornes, S.E. Hoelsbrekken, J.C. Ryan, S. Fossum, E. Dissen, Molecular cloning of KLRI1 and KLRI2, a novel pair of lectin-like natural killer-cell receptors with opposing signalling motifs, Immunogenetics 56(11) (2005) 833–9.

[46] A. Kupz, T.A. Scott, G.T. Belz, D.M. Andrews, M. Greyer, A.M. Lew, A.G. Brooks, M.J. Smyth, R. Curtiss, 3rd, S. Bedoui, R.A. Strugnell, Contribution of Thy1+ NK cells to protective IFN-γ production during Salmonella typhimurium infections, Proc Natl Acad Sci U S A 110(6) (2013) 2252-7.

[47] P.C. Saether, I.H. Westgaard, S.E. Hoelsbrekken, J. Benjamin, L.L. Lanier, S. Fossum, E. Dissen, KLRE/I1 and KLRE/I2: a novel pair of heterodimeric receptors that inversely regulate NK cell cytotoxicity, J Immunol 181(5) (2008) 3177–82.

[48] S.J. van der Stegen, D.M. Davies, S. Wilkie, J. Foster, J.K. Sosabowski, J. Burnet, L.M. Whilding, R.M. Petrovic, S. Ghaem-Maghami, S. Mather, J.P. Jeannon, A.C. Parente-Pereira, J. Maher, Preclinical in vivo modeling of cytokine release syndrome induced by ErbB-retargeted human T cells: identifying a window of therapeutic opportunity?, J Immunol 191(9) (2013) 4589–98.

[49] Y. Guo, W. Mao, L. Jin, L. Xia, J. Huang, X. Liu, P. Ni, Q. Shou, H. Fu, Flavonoid Group of Smilax glabra Roxb. Regulates the Anti-Tumor Immune Response Through the STAT3/HIF-1 Signaling Pathway, Front Pharmacol 13 (2022) 918975.

[50] W.F. Hawse, R.T. Cattley, T cells transduce T-cell receptor signal strength by generating different phosphatidylinositols, J Biol Chem 294(13) (2019) 4793–4805.

[51] B. Jin, Y. Liang, Y. Liu, L.X. Zhang, F.Y. Xi, W.J. Wu, Y. Li, G.H. Liu, Notch signaling pathway regulates T cell dysfunction in septic patients, Int Immunopharmacol 76 (2019) 105907.

[52] H. Tan, J. Ye, X. Luo, S. Chen, Q. Yin, L. Yang, Y. Li, Clonal expanded TRA and TRB subfamily T cells in peripheral blood from patients with diffuse large B-cell lymphoma, Hematology 15(2) (2010) 81–7.

[53] S.M. Hedrick, R. Hess Michelini, A.L. Doedens, A.W. Goldrath, E.L. Stone, FOXO transcription factors throughout T cell biology, Nat Rev Immunol 12(9) (2012) 649–61.

[54] S. Li, J. Ding, Y. Wang, X. Wang, L. Lv, CD155/TIGIT signaling regulates the effector function of tumor-infiltrating CD8+ T cell by NF-κB pathway in colorectal cancer, J Gastroenterol Hepatol 37(1) (2022) 154–163.

[55] F. Ding, X. Luo, Y. Tu, X. Duan, J. Liu, L. Jia, P. Zheng, Alpk1 Sensitizes Pancreatic Beta Cells to Cytokine-Induced Apoptosis via Upregulating TNF-α Signaling Pathway, Front Immunol 12 (2021) 705751.

[56] D.R. Sen, J. Kaminski, R.A. Barnitz, M. Kurachi, U. Gerdemann, K.B. Yates, H.W. Tsao, J. Godec, M.W. LaFleur, F.D. Brown, P. Tonnerre, R.T. Chung, D.C. Tully, T.M. Allen, N. Frahm, G.M. Lauer, E.J. Wherry, N. Yosef, W.N. Haining, The epigenetic landscape of T cell exhaustion, Science 354(6316) (2016) 1165-1169.

[57] E.J. Wherry, M. Kurachi, Molecular and cellular insights into T cell exhaustion, Nat Rev Immunol 15(8) (2015) 486–99.

[58] A. Tanaka, S. Sakaguchi, Regulatory T cells in cancer immunotherapy, Cell Res 27(1) (2017) 109–118.

[59] C. Tay, A. Tanaka, S. Sakaguchi, Tumor-infiltrating regulatory T cells as targets of cancer immunotherapy, Cancer Cell 41(3) (2023) 450–465.

[60] K.L. Banta, X. Xu, A.S. Chitre, A. Au-Yeung, C. Takahashi, W.E. O’Gorman, T.D. Wu, S. Mittman, R. Cubas, L. Comps-Agrar, A. Fulzele, E.J. Bennett, J.L. Grogan, E. Hui, E.Y. Chiang, I. Mellman, Mechanistic convergence of the TIGIT and PD-1 inhibitory pathways necessitates co-blockade to optimize anti-tumor CD8(+) T cell responses, Immunity 55(3) (2022) 512–526.e9.

[61] R.M. Gorczynski, IL-17 Signaling in the Tumor Microenvironment, Adv Exp Med Biol 1240 (2020) 47–58.

[62] M.T. Orr, L.L. Lanier, Inhibitory Ly49 receptors on mouse natural killer cells, Curr Top Microbiol Immunol 350 (2011) 67–87.

[63] M. Wang, H. Tao, P. Huang, Clinical significance of LARGE1 in progression of liver cancer and the underlying mechanism, Gene 779 (2021) 145493.

[64] Z.W. Liu, Z.J. Guo, A.L. Chu, Y. Zhang, B. Liang, X. Guo, T. Chai, R. Song, G. Hou, J.J. Yuan, High incidence of coding gene mutations in mitochondrial DNA in esophageal cancer, Mol Med Rep 16(6) (2017) 8537–8541.

[65] P. Pandey, B. Sliker, H.L. Peters, A. Tuli, J. Herskovitz, K. Smits, A. Purohit, R.K. Singh, J. Dong, S.K. Batra, D.W. Coulter, J.C. Solheim, Amyloid precursor protein and amyloid precursor-like protein 2 in cancer, Oncotarget 7(15) (2016) 19430–44.

[66] H. Zeng, L. Tang, CRIM1, the antagonist of BMPs, is a potential risk factor of cancer, Curr Cancer Drug Targets 14(7) (2014) 652–8.

[67] H. Kang, J. Fichna, K. Matlawska-Wasowska, D. Jacenik, The Expression Pattern of Adhesion G Protein-Coupled Receptor F5 Is Related to Cell Adhesion and Metastatic Pathways in Colorectal Cancer-Comprehensive Study Based on In Silico Analysis, Cells 11(23) (2022).

[68] K. Lawrenson, B. Grun, N. Lee, P. Mhawech-Fauceglia, J. Kan, S. Swenson, Y.G. Lin, T. Pejovic, J. Millstein, S.A. Gayther, NPPB is a novel candidate biomarker expressed by cancer-associated fibroblasts in epithelial ovarian cancer, Int J Cancer 136(6) (2015) 1390–401.

[69] T.Y. Bowley, I.V. Lagutina, C. Francis, S. Sivakumar, R.G. Selwyn, E. Taylor, Y. Guo, B.N. Fahy, B. Tawfik, D. Marchetti, The RPL/RPS gene signature of melanoma CTCs associates with brain metastasis, Cancer Res Commun 2(11) (2022) 1436–1448.

[70] T.Y. Bowley, S.D. Merkley, I.V. Lagutina, M.C. Ortiz, M. Lee, B. Tawfik, D. Marchetti, Targeting Translation and the Cell Cycle Inversely Affects CTC Metabolism but Not Metastasis, Cancers (Basel) 15(21) (2023).

[71] M. Yang, C.Y. Zhang, Interleukins in liver disease treatment, World J Hepatol 16(2) (2024) 140–145.

[72] K. Tuomela, A.R. Ambrose, D.M. Davis, Escaping Death: How Cancer Cells and Infected Cells Resist Cell-Mediated Cytotoxicity, Front Immunol 13 (2022) 867098.

[73] S. Kuerten, T.M. Nowacki, T.O. Kleen, R.J. Asaad, P.V. Lehmann, M. Tary-Lehmann, Dissociated production of perforin, granzyme B, and IFN-gamma by HIV-specific CD8(+) cells in HIV infection, AIDS Res Hum Retroviruses 24(1) (2008) 62–71.

[74] G. Hodge, J. Barnawi, C. Jurisevic, D. Moffat, M. Holmes, P.N. Reynolds, H. Jersmann, S. Hodge, Lung cancer is associated with decreased expression of perforin, granzyme B and interferon (IFN)-γ by infiltrating lung tissue T cells, natural killer (NK) T-like and NK cells, Clin Exp Immunol 178(1) (2014) 79–85.

[75] R.G. Urdinguio, A.F. Fernandez, A. Moncada-Pazos, C. Huidobro, R.M. Rodriguez, C. Ferrero, P. Martinez-Camblor, A.J. Obaya, T. Bernal, A. Parra-Blanco, L. Rodrigo, M. Santacana, X. Matias-Guiu, B. Soldevilla, G. Dominguez, F. Bonilla, S. Cal, C. Lopez-Otin, M.F. Fraga, Immune-dependent and independent antitumor activity of GM-CSF aberrantly expressed by mouse and human colorectal tumors, Cancer Res 73(1) (2013) 395–405.

[76] V. Bretscher, D. Andreutti, P. Neuville, M. Martin, F. Martin, O. Lefebvre, C. Gilles, G. Benzonana, G. Gabbiani, GM-CSF expression by tumor cells correlates with aggressivity and with stroma reaction formation, J Submicrosc Cytol Pathol 32(4) (2000) 525–33.

[77] I.S. Hong, Stimulatory versus suppressive effects of GM-CSF on tumor progression in multiple cancer types, Exp Mol Med 48(7) (2016) e242.

[78] M. Thorn, P. Guha, M. Cunetta, N.J. Espat, G. Miller, R.P. Junghans, S.C. Katz, Tumor-associated GM-CSF overexpression induces immunoinhibitory molecules via STAT3 in myeloid-suppressor cells infiltrating liver metastases, Cancer Gene Ther 23(6) (2016) 188–98.

[79] M.J. Sweet, D.A. Hume, CSF-1 as a regulator of macrophage activation and immune responses, Arch Immunol Ther Exp (Warsz) 51(3) (2003) 169–77.

[80] D.A. Hume, K.P. MacDonald, Therapeutic applications of macrophage colony-stimulating factor-1 (CSF-1) and antagonists of CSF-1 receptor (CSF-1R) signaling, Blood 119(8) (2012) 1810–20.

[81] J. Zhou, C. Tulotta, P.D. Ottewell, IL-1β in breast cancer bone metastasis, Expert Rev Mol Med 24 (2022) e11.

[82] C. Rébé, F. Ghiringhelli, Interleukin-1β and Cancer, Cancers (Basel) 12(7) (2020).

[83] M. Mizui, Natural and modified IL-2 for the treatment of cancer and autoimmune diseases, Clin Immunol 206 (2019) 63–70.

[84] L.H. Corrêa, R. Corrêa, C.M. Farinasso, L.P. de Sant’Ana Dourado, K.G. Magalhães, Adipocytes and Macrophages Interplay in the Orchestration of Tumor Microenvironment: New Implications in Cancer Progression, Front Immunol 8 (2017) 1129.

[85] J.M. Whitworth, R.D. Alvarez, Evaluating the role of IL-12 based therapies in ovarian cancer: a review of the literature, Expert Opin Biol Ther 11(6) (2011) 751–62.

[86] N. Kumar, A. Vyas, S.K. Agnihotri, N. Chattopadhyay, M. Sachdev, Small secretory proteins of immune cells can modulate gynecological cancers, Semin Cancer Biol 86(Pt 3) (2022) 513–531.

[87] G.M. Konjević, A.M. Vuletić, K.M. Mirjačić Martinović, A.K. Larsen, V.B. Jurišić, The role of cytokines in the regulation of NK cells in the tumor environment, Cytokine 117 (2019) 30–40.

[88] T.A. Wynn, IL-13 effector functions, Annu Rev Immunol 21 (2003) 425–56.

[89] S. Chen, J. Jiao, D. Jiang, Z. Wan, L. Li, K. Li, L. Xu, Z. Zhou, W. Xu, J. Xiao, T-box transcription factor Brachyury in lung cancer cells inhibits macrophage infiltration by suppressing CCL2 and CCL4 chemokines, Tumour Biol 36(8) (2015) 5881–90.

[90] C. Ottaviani, F. Nasorri, C. Bedini, O. de Pità, G. Girolomoni, A. Cavani, CD56brightCD16(-) NK cells accumulate in psoriatic skin in response to CXCL10 and CCL5 and exacerbate skin inflammation, Eur J Immunol 36(1) (2006) 118–28.

[91] S. Ghafouri-Fard, M. Shahir, M. Taheri, A. Salimi, A review on the role of chemokines in the pathogenesis of systemic lupus erythematosus, Cytokine 146 (2021) 155640.

[92] M. Gouwy, M. Schiraldi, S. Struyf, J. Van Damme, M. Uguccioni, Possible mechanisms involved in chemokine synergy fine tuning the inflammatory response, Immunol Lett 145(1-2) (2012) 10–4.

[93] R. Janssens, S. Struyf, P. Proost, The unique structural and functional features of CXCL12, Cell Mol Immunol 15(4) (2018) 299–311.

[94] R. Janssens, S. Struyf, P. Proost, Pathological roles of the homeostatic chemokine CXCL12, Cytokine Growth Factor Rev 44 (2018) 51–68.

[95] P. Dent, A. Yacoub, J. Contessa, R. Caron, G. Amorino, K. Valerie, M.P. Hagan, S. Grant, R. Schmidt-Ullrich, Stress and radiation-induced activation of multiple intracellular signaling pathways, Radiat Res 159(3) (2003) 283–300.

[96] S.R. Burks, R.M. Lorsung, M.E. Nagle, T.W. Tu, J.A. Frank, Focused ultrasound activates voltage-gated calcium channels through depolarizing TRPC1 sodium currents in kidney and skeletal muscle, Theranostics 9(19) (2019) 5517–5531.

[97] T. Benavides Damm, M. Egli, Calcium’s role in mechanotransduction during muscle development, Cell Physiol Biochem 33(2) (2014) 249–72.

[98] E. Memari, D. Khan, R. Alkins, B. Helfield, Focused ultrasound-assisted delivery of immunomodulating agents in brain cancer, J Control Release 367 (2024) 283–299.

[99] F. Ivanovski, M. Meško, T. Lebar, M. Rupnik, D. Lainšček, M. Gradišek, R. Jerala, M. Benčina, Ultrasound-mediated spatial and temporal control of engineered cells in vivo, Nat Commun 15(1) (2024) 7369.

[100] B.A. Runyon, Care of patients with ascites, N Engl J Med 330(5) (1994) 337–42.

[101] S.L. Parsons, S.A. Watson, R.J. Steele, Malignant ascites, Br J Surg 83(1) (1996) 6–14.

[102] H. Woopen, J. Sehouli, Current and future options in the treatment of malignant ascites in ovarian cancer, Anticancer Res 29(8) (2009) 3353–9.

[103] X. Han, L. An, D. Yan, M. Hiroshi, W. Ding, M. Zhang, G. Xu, Y. Sun, G. Yuan, M. Wang, N. Zhao, J. Sun, X. Zhu, P. Du, Combined antitumor effects of P-5m octapeptide and 5-fluorouracil on a murine model of H22 hepatoma ascites, Exp Ther Med 16(3) (2018) 1586–1592.

[104] I. Masuda, J. Hirose, Animal models of pathologic calcification, Curr Opin Rheumatol 14(3) (2002) 287–91.

[105] I.T. Olszak, M.C. Poznansky, R.H. Evans, D. Olson, C. Kos, M.R. Pollak, E.M. Brown, D.T. Scadden, Extracellular calcium elicits a chemokinetic response from monocytes in vitro and in vivo, J Clin Invest 105(9) (2000) 1299–305.

[106] P.R. Dominguez-Gutierrez, S. Kusmartsev, B.K. Canales, S.R. Khan, Calcium Oxalate Differentiates Human Monocytes Into Inflammatory M1 Macrophages, Front Immunol 9 (2018) 1863.

[107] S. Kusmartsev, P.R. Dominguez-Gutierrez, B.K. Canales, V.G. Bird, J. Vieweg, S.R. Khan, Calcium Oxalate Stone Fragment and Crystal Phagocytosis by Human Macrophages, J Urol 195(4 Pt 1) (2016) 1143-51.

[108] R. de Water, P.J. Leenen, C. Noordermeer, A.L. Nigg, A.B. Houtsmuller, D.J. Kok, F.H. Schröder, Cytokine production induced by binding and processing of calcium oxalate crystals in cultured macrophages, Am J Kidney Dis 38(2) (2001) 331–8.

[109] H. Alshetaiwi, N. Pervolarakis, L.L. McIntyre, D. Ma, Q. Nguyen, J.A. Rath, K. Nee, G. Hernandez, K. Evans, L. Torosian, A. Silva, C. Walsh, K. Kessenbrock, Defining the emergence of myeloid-derived suppressor cells in breast cancer using single-cell transcriptomics, Sci Immunol 5(44) (2020).

[110] V. Bronte, S. Brandau, S.H. Chen, M.P. Colombo, A.B. Frey, T.F. Greten, S. Mandruzzato, P.J. Murray, A. Ochoa, S. Ostrand-Rosenberg, P.C. Rodriguez, A. Sica, V. Umansky, R.H. Vonderheide, D.I. Gabrilovich, Recommendations for myeloid-derived suppressor cell nomenclature and characterization standards, Nat Commun 7 (2016) 12150.

[111] H. Lewinsky, E.G. Gunes, K. David, L. Radomir, M.P. Kramer, B. Pellegrino, M. Perpinial, J. Chen, T.F. He, A.G. Mansour, K.Y. Teng, S. Bhattacharya, E. Caserta, E. Troadec, P. Lee, M. Feng, J. Keats, A. Krishnan, M. Rosenzweig, J. Yu, M.A. Caligiuri, Y. Cohen, O. Shevetz, S. Becker-Herman, F. Pichiorri, S. Rosen, I. Shachar, CD84 is a regulator of the immunosuppressive microenvironment in multiple myeloma, JCI Insight 6(4) (2021).

[112] A. Marom, A.F. Barak, M.P. Kramer, H. Lewinsky, I. Binsky-Ehrenreich, S. Cohen, A. Tsitsou-Kampeli, V. Kalchenko, Y. Kuznetsov, V. Mirkin, N. Dezorella, M. Shapiro, P.L. Schwartzberg, Y. Cohen, L. Shvidel, M. Haran, S. Becker-Herman, Y. Herishanu, I. Shachar, CD84 mediates CLL-microenvironment interactions, Oncogene 36(5) (2017) 628–638.

[113] Y. Si, S.F. Merz, P. Jansen, B. Wang, K. Bruderek, P. Altenhoff, S. Mattheis, S. Lang, M. Gunzer, J. Klode, A. Squire, S. Brandau, Multidimensional imaging provides evidence for down-regulation of T cell effector function by MDSC in human cancer tissue, Sci Immunol 4(40) (2019).

[114] M.D. Cahalan, K.G. Chandy, The functional network of ion channels in T lymphocytes, Immunol Rev 231(1) (2009) 59–87.

[115] S. Feske, H. Wulff, E.Y. Skolnik, Ion channels in innate and adaptive immunity, Annu Rev Immunol 33 (2015) 291–353.

[116] M. Vaeth, S. Kahlfuss, S. Feske, CRAC Channels and Calcium Signaling in T Cell-Mediated Immunity, Trends Immunol 41(10) (2020) 878–901.

[117] Y.J. Park, S.A. Yoo, M. Kim, W.U. Kim, The Role of Calcium-Calcineurin-NFAT Signaling Pathway in Health and Autoimmune Diseases, Front Immunol 11 (2020) 195.

[118] S. Feske, E.Y. Skolnik, M. Prakriya, Ion channels and transporters in lymphocyte function and immunity, Nat Rev Immunol 12(7) (2012) 532–47.

[119] X. Meng, X. Wu, Y. Zheng, K. Shang, R. Jing, P. Jiao, C. Zhou, J. Zhou, J. Sun, Exploiting Ca(2+) signaling in T cells to advance cancer immunotherapy, Semin Immunol 49 (2020) 101434.

[120] E.C. Schwarz, B. Qu, M. Hoth, Calcium, cancer and killing: the role of calcium in killing cancer cells by cytotoxic T lymphocytes and natural killer cells, Biochim Biophys Acta 1833(7) (2013) 1603–11.

[121] A. Maul-Pavicic, S.C. Chiang, A. Rensing-Ehl, B. Jessen, C. Fauriat, S.M. Wood, S. Sjöqvist, M. Hufnagel, I. Schulze, T. Bass, W.W. Schamel, S. Fuchs, H. Pircher, C.A. McCarl, K. Mikoshiba, K. Schwarz, S. Feske, Y.T. Bryceson, S. Ehl, ORAI1-mediated calcium influx is required for human cytotoxic lymphocyte degranulation and target cell lysis, Proc Natl Acad Sci U S A 108(8) (2011) 3324–9.

[122] L. Kaschek, S. Zöphel, A. Knörck, M. Hoth, A calcium optimum for cytotoxic T lymphocyte and natural killer cell cytotoxicity, Semin Cell Dev Biol 115 (2021) 10–18.

[123] Y. Wang, Z. Liu, Y. Qi, J. Wu, B. Liu, X. Cui, Activin A, a Novel Chemokine, Induces Mouse NK Cell Migration via AKT and Calcium Signaling, Cells 13(9) (2024).

[124] A.M. Shenoy, R.A. Sidner, Z. Brahmi, Signal transduction in cytotoxic lymphocytes: decreased calcium influx in NK cell inactivated with sensitive target cells, Cell Immunol 147(2) (1993) 294–301.

[125] X. Zhou, K.S. Friedmann, H. Lyrmann, Y. Zhou, R. Schoppmeyer, A. Knörck, S. Mang, C. Hoxha, A. Angenendt, C.S. Backes, C. Mangerich, R. Zhao, S. Cappello, G. Schwär, C. Hässig, M. Neef, B. Bufe, F. Zufall, K. Kruse, B.A. Niemeyer, A. Lis, B. Qu, C. Kummerow, E.C. Schwarz, M. Hoth, A calcium optimum for cytotoxic T lymphocyte and natural killer cell cytotoxicity, J Physiol 596(14) (2018) 2681–2698.

[126] G. Luo, X. Li, J. Lin, G. Ge, J. Fang, W. Song, G.G. Xiao, B. Zhang, X. Peng, Y. Duo, B.Z. Tang, Multifunctional Calcium-Manganese Nanomodulator Provides Antitumor Treatment and Improved Immunotherapy via Reprogramming of the Tumor Microenvironment, ACS Nano 17(16) (2023) 15449–15465.

[127] J. An, K. Zhang, B. Wang, S. Wu, Y. Wang, H. Zhang, Z. Zhang, J. Liu, J. Shi, Nanoenabled Disruption of Multiple Barriers in Antigen Cross-Presentation of Dendritic Cells via Calcium Interference for Enhanced Chemo-Immunotherapy, ACS Nano 14(6) (2020) 7639–7650.

[128] Y. Lin, X. Wang, X. Huang, J. Zhang, N. Xia, Q. Zhao, Calcium phosphate nanoparticles as a new generation vaccine adjuvant, Expert Rev Vaccines 16(9) (2017) 895–906.

[129] P. Zhou, D. Xia, Z. Ni, T. Ou, Y. Wang, H. Zhang, L. Mao, K. Lin, S. Xu, J. Liu, Calcium silicate bioactive ceramics induce osteogenesis through oncostatin M, Bioact Mater 6(3) (2021) 810–822.

[130] A. Yamaguchi, N. Maeshige, H. Noguchi, J. Yan, X. Ma, M. Uemura, D. Su, H. Kondo, K. Sarosiek, H. Fujino, Pulsed ultrasound promotes secretion of anti-inflammatory extracellular vesicles from skeletal myotubes via elevation of intracellular calcium level, Elife 12 (2023).

[131] R.E. Kumon, M. Aehle, D. Sabens, P. Parikh, D. Kourennyi, C.X. Deng, Ultrasound-induced calcium oscillations and waves in Chinese hamster ovary cells in the presence of microbubbles, Biophys J 93(6) (2007) L29–31.

[132] C.W. Yoon, N.S. Lee, K.M. Koo, S. Moon, K. Goo, H. Jung, C. Yoon, H.G. Lim, K.K. Shung, Investigation of Ultrasound-Mediated Intracellular Ca(2+) Oscillations in HIT-T15 Pancreatic β-Cell Line, Cells 9(5) (2020).

[133] S. Zanos, D. Ntiloudi, J. Pellerito, R. Ramdeo, J. Graf, K. Wallace, V. Cotero, J. Ashe, J. Moon, M. Addorisio, D. Shoudy, T.R. Coleman, M. Brines, C. Puleo, K.J. Tracey, S.S. Chavan, Focused ultrasound neuromodulation of the spleen activates an anti-inflammatory response in humans, Brain Stimul 16(3) (2023) 703–711.

[134] Y.C. Chu, J. Lim, A. Chien, C.C. Chen, J.L. Wang, Activation of Mechanosensitive Ion Channels by Ultrasound, Ultrasound Med Biol 48(10) (2022) 1981–1994.

[135] J. Zhu, Q. Xian, X. Hou, K.F. Wong, T. Zhu, Z. Chen, D. He, S. Kala, S. Murugappan, J. Jing, Y. Wu, X. Zhao, D. Li, J. Guo, Z. Qiu, L. Sun, The mechanosensitive ion channel Piezo1 contributes to ultrasound neuromodulation, Proc Natl Acad Sci U S A 120(18) (2023) e2300291120.

[136] D. Li, Y. Yong, C. Qiao, H. Jiang, J. Lin, J. Wei, Y. Zhou, F. Li, Low-Intensity Pulsed Ultrasound Dynamically Modulates the Migration of BV2 Microglia, Ultrasound Med Biol 51(3) (2025) 494–507.

[137] X. Hou, J. Jing, Y. Jiang, X. Huang, Q. Xian, T. Lei, J. Zhu, K.F. Wong, X. Zhao, M. Su, D. Li, L. Liu, Z. Qiu, L. Sun, Nanobubble-actuated ultrasound neuromodulation for selectively shaping behavior in mice, Nat Commun 15(1) (2024) 2253.

[138] Z. Fan, R.E. Kumon, J. Park, C.X. Deng, Intracellular delivery and calcium transients generated in sonoporation facilitated by microbubbles, J Control Release 142(1) (2010) 31–9.

[139] A. Tsukamoto, S. Higashiyama, K. Yoshida, Y. Watanabe, K.S. Furukawa, T. Ushida, Stable cavitation induces increased cytoplasmic calcium in L929 fibroblasts exposed to 1-MHz pulsed ultrasound, Ultrasonics 51(8) (2011) 982–90.

[140] W. Dong, G. Wang, Y. Chai, W. Li, S. Liu, H. Liu, W. Guo, S. Li, X. He, M. Wan, Z. Li, Y. Zong, Two-step ultrasonic cavitation controlled delivery of brain exogenous nucleic acids for ischemic stroke using acoustic-cationic-polymeric-nanodroplets, Drug Deliv Transl Res (2025).

[141] D. Przystupski, D. Baczyńska, J. Rossowska, J. Kulbacka, M. Ussowicz, Calcium ion delivery by microbubble-assisted sonoporation stimulates cell death in human gastrointestinal cancer cells, Biomed Pharmacother 179 (2024) 117339.

[142] R. Zhang, R. El-Mayta, T.J. Murdoch, C.C. Warzecha, M.M. Billingsley, S.J. Shepherd, N. Gong, L. Wang, J.M. Wilson, D. Lee, M.J. Mitchell, Helper lipid structure influences protein adsorption and delivery of lipid nanoparticles to spleen and liver, Biomater Sci 9(4) (2021) 1449–1463.

[143] I.A. Khalil, M.A. Younis, S. Kimura, H. Harashima, Lipid Nanoparticles for Cell-Specific in Vivo Targeted Delivery of Nucleic Acids, Biol Pharm Bull 43(4) (2020) 584–595.

[144] S. Kimura, I.A. Khalil, Y.H.A. Elewa, H. Harashima, Novel lipid combination for delivery of plasmid DNA to immune cells in the spleen, J Control Release 330 (2021) 753–764.

[145] T. Chaiwarit, S.R. Sommano, P. Rachtanapun, N. Kantrong, W. Ruksiriwanich, M. Kumpugdee-Vollrath, P. Jantrawut, Development of Carboxymethyl Chitosan Nanoparticles Prepared by Ultrasound-Assisted Technique for a Clindamycin HCl Carrier, Polymers (Basel) 14(9) (2022).

[146] Z. Cao, X. Yang, W. Yang, F. Chen, W. Jiang, S. Zhan, F. Jiang, J. Li, C. Ye, L. Lang, S. Zhang, Z. Feng, X. Lai, Y. Liu, L. Mao, H. Cai, Y. Teng, J. Xie, Modulation of Dendritic Cell Function via Nanoparticle-Induced Cytosolic Calcium Changes, ACS Nano 18(10) (2024) 7618–7632.

